# Cocaine-dependent acquisition of locomotor sensitization and conditioned place preference requires D1 dopaminergic signaling through a cyclic AMP, NCS-Rapgef2, ERK and Egr-1/Zif268 pathway

**DOI:** 10.1101/2020.07.02.184754

**Authors:** Sunny Zhihong Jiang, Sean Sweat, Sam Dahlke, Kathleen Loane, Gunner Drossel, Wenqin Xu, Hugo A. Tejeda, Charles R. Gerfen, Lee E. Eiden

## Abstract

Elucidation of the underlying mechanism of dopamine signaling to ERK that underlies plasticity in dopamine D1 receptor expressingneurons leadingto acquired cocaine preference is incomplete. NCS-Rapgef2 is a novel cAMP effector, expressed in neuronal and endocrine cells in adult mammals, that is required for D1 dopamine receptor-dependent ERK phosphorylation in mouse brain. In this report, we studied the effects of abrogating NCS-Rapgef2 expression on cAMP-dependent ERK→Egr-1/zif268 signaling in cultured neuroendocrine cells; in D1 medium spiny neurons (MSNs) of nucleus accumbens slices; and in mouse brain in a region-specific manner. NCS-Rapgef2 gene deletion in the nucleus accumbens (NAc) in adult mice, using AAV-mediated expression of cre recombinase, eliminated cocaine-induced ERK phosphorylation and Egr-1/Zif268 upregulation in D1-MSNs and cocaine-induced behaviors including locomotor sensitization and conditioned place preference (CPP). Abrogation of NCS-Rapgef2 gene expression in medium prefrontal cortex and basolateral amygdala, by crossing mice bearing a floxed Rapgef2 allele with a cre mouse line driven by calcium/calmodulin-dependent kinase IIα promoter also eliminated cocaine-induced phospho-ERK activation and Egr-1/Zif268 induction, but without effect on the cocaine-induced behaviors. Our results indicate that NCS-Rapgef2 signaling to ERK in dopamine D1-receptor expressing neurons in the NAc, butnotin corticolimbic areas, contributes to cocaine-induced locomotor sensitization and CPP. Ablation of cocaine-dependent ERK activation by elimination of NCS-Rapgef2 occurred with no effect on phosphorylation of CREB in D1 dopaminoceptive neurons of NAc. This study reveals a new cAMP-dependent signaling pathway for cocaine-induced behavioral adaptations, mediated through NCS-Rapgef2/phospho-ERK activation, independently of PKA/CREB signaling.

**SIGNIFICANCE STATEMENT:** ERK phosphorylation in dopamine D1 receptor expressing neurons exerts a pivotal role in psychostimulant-induced neuronal gene regulation and behavioral adaptation, including locomotor sensitization and drug preference in rodents. In this study, we examined the role of dopamine signaling through the D1 receptor via a novel pathway initiated through the cAMP-activated guanine nucleotide exchange factor NCS-Rapgef2 in mice. NCS-Rapgef2 in the nucleus accumbens is required for activation of ERK and Egr-1/Zif268 in D1 dopaminoceptive neurons after acute cocaine administration, and subsequentenhanced locomotor response anddrugseeking behavior after repeated cocaine administration. This novel component in dopamine signaling provides a potential new target for intervention in psychostimulant-shaped behaviors, and new understanding of how D1-MSNs encode the experience of psychomotor stimulant exposure.

## INTRODUCTION

Behavioral plasticity, including locomotor sensitization and development of drug preference, is associated with psychomotor stimulant-induced neuroadaptation in brain reward circuits linked to addiction (Robinson and Berridge, 2000; Laakso et al., 2002; Fernando and Robbins, 2011; Uhl et al., 2019). Neurons in striatum, prefrontal cortex, amygdala and hippocampus that receive dopaminergic input from the ventral tegmental area (VTA) are central to psychomotor stimulant-induced neuroadaptation (Rebec and Sun, 2005; Nestler and Luscher, 2019). Amphetamine and cocaine increase the concentration of dopamine released from nerve terminals in these brain regions (Heien et al., 2005; Nestler and Luscher, 2019). Dopamine acts at receptors promoting either an increase (D1 type) or a decrease (D2 type) in cyclic AMP production in distinct dopaminoceptive neurons. A balance between these two processes contributes to the behavioral sequelae, including addiction, of repeated psychomotor stimulant use (Lobo and Nestler, 2011). Cellular mechanisms for the behavioral effects of psychomotor stimulant exposure have been soughtby examining dopamine signalingthrough the Gs-coupled D1 receptor in various brain areas. One of these is the nucleus accumbens (NAc), which contains GABAergic medium spiny neurons (MSNs) neurons whose activity is modulated by dopamine, and represent the afferent targets of the reward pathway from VTA to NAc in mammalian brain (Hikida et al., 2010; Russo and Nestler, 2013).

The MAP kinase ERK is activated *in vivo* upon treatment with D1 agonists, cocaine or amphetamine (Valjent et al., 2000; Gutierrez-Arenas et al., 2014; Jin et al., 2019) in brain regions innervated by dopaminergic neurons. Pharmacological and genetic manipulations have further implicated ERK activation in transcriptional effects of psychomotor stimulants, notably expression of the immediate-early gene egr1/zif268, linked to psychomotor stimulant-induced behaviors (Valjent et al., 2006c). However, genetic and pharmacological tools have limited cellular and anatomical specificity for analysis of these pathways. For example, generalized effects on locomotor activity and co-regulatory effects of ERK1 and ERK2 complicate interpretation of ERK1/2 knockout effects on both behavior and cell signaling (Mazzucchelli et al., 2002; Ferguson et al., 2006; Hutton et al., 2017). Consequently, it has been difficult to identify a signaling pathway arising from D1 receptor activation leading to the specific ERK activation and Egr1/Zif268 induction that mediates cellular plasticity within dopaminoceptive neurons, in defined brain circuits, responsible for the altered behaviors typical of repeated psychomotor stimulant administration.

In particular, the role of protein kinase A (PKA) as the downstream effector of cyclic AMP in the D1-dependent activation of Egr-1/Zif268 by ERK (Funahashi et al., 2019), leading to cocaine-dependent locomotor sensitization, has been difficult to confirm. This suggests that a second cAMP-dependent pathway for ERK regulation of Egr-1/Zif268 induction, and subsequent behavioral effects of cocaine mediated through dopamine, may exist in D1-MSNs.

We have previously identified a novel pathway for ERK activation via the novel cAMP effector NCS-Rapgef2 in NS-1 neuroendocrine cells in culture (Emery and Eiden, 2012; Emery et al., 2013; Emery et al., 2014; Emery et al., 2017a). NCS-Rapgef2 is activated by cAMP binding to facilitate GDP-GTP exchange on the Rap1 molecule, allowing Rap1 to further stimulate the MAP kinase cascade culminating in ERK phosphorylation/activation (see Jiang et al., 2017). Using this assay as a read-out for the NCS-Rapgef2 dependent cAMP pathway, we have shown that the dopamine D1 receptor couples to NCS-Rapgef2 to activate ERK in neuroendocrine cells, and further demonstrated that this pathway is required for D1-dependent activation of ERK in hippocampus and other forebrain regions of the adult mouse brain *in vivo* (Jiang et al., 2017).

These data open the possibility that a direct pathway for activation of ERK by the Gs/olf- coupled D1 receptor leading to Egr-1/Zif268 induction exists in D1 dopaminoceptive neurons involved in the behavioraleffects of repeated cocaine administration. Here, we have asked whether the NCS-Rapgef2-mediated activation of ERK after cyclic AMP elevation by the Gs-coupled D1 receptor is responsible for the behavioral effects of psychomotor stimulant administration, and if so, in which neuronal ensembles or circuits it might be relevant.

## METHODS AND MATERIALS

### Animals and drugs administered *in vivo*

Mice (wild-type or transgenic) on C57BL6J background were housed 2-5 per cage and acclimatized to 12-hour light/12-hour dark cycle with food and water *ad libitum*. Animal care, drug treatment and surgeries were approved by the NIMH Institutional Animal Care and Use Committee (ACUC) and conducted in accordance with the NIH guidelines. The floxed Rapgef2 mouse strain Rapgef2^cko/cko^ and corticolimbic Rapgef2 KO mouse strain Camk2α-cre^+/-^::Rapgef2^cko/cko^ were generated as described previously (Jiang et al., 2017). The transgenic Drd1a-tdTomato BAC mouse (The Jackson Laboratory, #016204), expressing tdTomato selectively in D1-dopaminoceptive neurons (Shuen et al., 2008), was obtained from the Jackson Laboratory. Pharmaceutical grade D1 dopamine receptor agonist SKF81297, D2 antagonist eticlopride, and cocaine were purchased from Sigma (St. Louis, MO) and dissolved in saline. All drugs used *in vivo* were administrated by intraperitoneal (*i.p.*) injection.

### Brain slice pharmacology

250 μm thick coronal slices of mouse brain containing the nucleus accumbens were prepared as described previously for electrophysiological examination under perfusion (Tejeda et al., 2017), but were instead incubated in a static bath of continuously aerated artificial cerebrospinal fluid, total volume 10 ml per slice, to which various drugs and drug combinations were added to examine the parameters of dopamine and cAMP-dependent signaling in dopaminoceptive cells of core and shell of this brain region. Inhibitors or vehicle were added for a 10 min period prior to addition of 10 μM SKF81297, or vehicle. Incubation at 34°C was continued for 30 min, and slices then quickly removed to cold 10% formalin in phosphate-buffered saline for immunohistochemical staining and microscopical analysis as described below. N6-phenyl, 9-tetrahydrofuranyl adenine (EL1101), a preferential inhibitor of the Rap guanine nucleotide exchange activity of NCS-Rapgef2 compared to cAMP effectors protein kinase A and Epac, was synthesized and purified as described previously (Emery et al., 2017b). All drugs were initially dissolved in DMSO, present to a final concentration of 0.1% in all experiments. Final concentration of U0126 was 10 μM, and final concentration of EL1101 was 100 μM, in slice experiments.

### Cell lines and cell culture

NS-1 cells (Emery et al., 2013) were cultured in Roswell Park Memorial Institute (RPMI) 1640 medium supplemented with 10% HyClone horse serum (GEHealthcare, Marlborough, MA), 5% heat-inactivated fetal bovine serum, 2 mM L-glutamine, 100 U/ml penicillin, and 100 µg/ml streptomycin. The established NS-1 cell line with NCS-Rapgef2 knocked-out was described previously (Jiang et al., 2017). The cell-permeable cyclic AMP analog 8-chlorophenylthio-cyclic AMP (8-CPT-cAMP) was purchased from Biolog Life Science Institute (Bremen, Germany) and the stock solution was prepared at 100 mM in DMSO, diluted in culture media to a final concentration of 100 μM.

### Viral injection

Modified AAV vectors were obtained from Penn Vector Core. Rapgef2 ^cko/cko^ mice were bilaterally injected with AAV9.hSynap.HI.eGFP-Cre.WPRE.SV40, encoding eGFP-fused Cre recombinase under the control of the human synapsin promoter, to knock out Rapgef2 expression in nucleus accumbens on both sides of the brain. AAV9.hSynap.eGFP.WPRE.SV40, encoding eGFP without Cre, also under the controlof the synapsinpromoter, was used as a control. Surgeries and viral injection were conducted according to the “NIH-ARAC guidelines for survival rodent surgery”. Briefly, animals were anesthetized, shaved and mounted into the stereotaxic apparatus. A small midline incision was then made and the pericranial tissue was teased away from the skull with an ethanol-soaked swab to enable identification of the Bregma and Lambda areas. Using a small hand-held drill, a very small hole was made in the skull according to the calculated coordinates for nucleus accumbens (Bregma 1.5 mm, ML 0.7 mm, DV 3.8 mm). A Hamilton syringe (pre-loaded with viral particles) was then slowly lowered, penetrating the dura to the determined depth. A volume of 0.3 μl of the virus (∼0.5×10^9^ infectious particle per microliter) was slowly (∼0.1 μl/min) injected into the brain. A 2∼3-minute wait is performed before the needle is very slowly retracted from the brain to prevent backflow of the viral vector. The animals were allowed to recover and subjected to behavioral tests four weeks after viral injection.Brain injection sites were histologically verified with counterstaining from post-mortem sections and plotted on standardized sections from the stereotaxic atlas.

### Immunohistochemistry

Immunohistochemistry was conducted as previously described (Jiang et al., 2017) after animal perfusion with 4% paraformaldehyde. Mouse brains were sectioned by Vibratome at a 40 μm thickness. Free-floating sections were washed in TBS containing 0.5% Triton X-100 (TBST; 3 washes, 15 min), incubated at room temperature in blocking solution (10% normal goat or donkey serum in PBST; 1h), and then incubated in primary antibody diluted in blocking solution overnight at 4°C. The following day sections were washed in TBST (3 washes, 15 min), incubated in the dark in Alexa 555-conjugated goat anti-rabbit-IgG, Alexa 488-conjugated goat anti-mouse IgG, Alexa 488-conjugated goat anti-rabbit-IgG, Alexa 555 -conjugated donkey anti-goat-IgG or Alexa 488-conjugated donkey anti-rabbit-IgG (1:300; Life Technologies, Grand Island, NY) for 2 h following primary antibody incubation. Sections were mounted in Vectashield (Vector Laboratories, Burlingame, CA). The primary antibodies used were rabbit anti-pERK (1:1500, Cell Signaling, Danvers, MA), anti-pCREB (1:1000, Cell Signaling, Denvers, MA), mouse anti-PSD95 (1:1000, NeuroMab, Davis, CA), mouse anti-Bassoon (1:250, Enzo Life Sciences, Inc, Farmingdale, NY), goat anti-tdTomato (1:1000, MyBiosource, Inc, San Diego, CA), rabbit anti-Egr1 (588) (1:7500, Santa Cruz, CA), rabbit anti-Rapgef2 (NNLE-2, custom-made by Anaspec, see ref. (Jiang et al., 2017). Confocal images were obtained on a Zeiss LSM 510 confocal microscope at the National Institute of Neurological Disorders and Stroke Light Imaging Facility. Immunoreactive (IR) signals of NCS-Rapgef2, phospho-ERK and Egr-1 from differentbrain areas as indicated were quantified by NIH Image J using the mean gray values of integrated density after being converted to gray scale. To quantify the NCS-Rapgef2, phospho-ERK and phospho-CREB IR signals in AAV viral targeted neurons in the NAc, confocal images with double fluorescence channels were imported into Fiji ImageJ and split into different channels. Images in GFP channel were inverted and made into binary mask after adjusting threshold. GFP positive neurons were selected as ROI following particle analysis. The ROI selections were overlaid to the NCS-Rapgef2, phospho-ERK or phospho-CREB channel. Cell number for phospho-ERK positive neurons or mean gray value for NCS-Rapgef2, phospho-CREB IR in ROIs created from GFPpositive neurons were measured. GFP signal from AAV-hSyn-eGFP control viral injection and AAV-hSyn-cre-eGFP have different ROIs since cre-eGFP is localized to nuclei and eGFP distributes throughout the cell. pCREB signal is localized to nuclei, so ROI selections from cre-eGFP and eGFP can both completely cover pCREB signal for GFP+/pCREB+ double positive neurons, while this is not the case for GPF+/phospho-ERK (pERK) double positive neurons. Since all neurons show basal pCREB expression, quantification of pCREB is more accurate using intensity. However, pERK signal distributes throughout the cell, so that ROI selections from cre-eGFP do not completely cover pERK signal for the GFP+/pERK+ double positive neurons. Thus, more accurate quantification of pERK/GPF co-expression was more accurately assessed via counting of cell number.

### Western blot for cultured cell and mouse tissues

NS-1 cells were seeded in collagen 1-coated 12-well plates and collected in ice-cold RIPA lysis buffer (Thermo Scientific, Rockford, IL) containing 1x Halt™ Protease and Phosphatase Inhibitor Cocktail (Thermo Scientific). Tissues were punched from 0.5 mm coronal sections of mouse brains, weighed, and snap-frozen. Samples were sonicated on ice with RIPA buffer supplemented with Halt Protease and Phosphatase Inhibitor Cocktail (Thermo Scientific). RIPA-insoluble fractions were removed by centrifugation (3000 rpm for 10 min at 4°C). Supernatants were retained, and protein concentration was determined by MicroBCA Protein Assay kit (Thermo Scientific) according to the protocol provided by the manufacturer. Western blot was performed as described previously (Jiang et al., 2017). The rabbit monoclonal antibodies including anti-ERK (Cell Signaling Technology, Cat# 9102, 1:1000), rabbit anti-phospho-ERK (Cell Signaling Technology, Cat# 9101, 1:1000), rabbit anti-Egr-1 (15F7)(Cell Signaling Technology, Cat# 4153; 1:1000) and anti-GAPDH (D16H11) (Cell Signaling Technology, Cat# 5174; 1:1000), were all purchased from Cell signaling Technology and used in the Western blot according to the manufacturer’s recommendations. Immuoreactive bands were visualized with a SuperSignal West Dura Chemiluminescent Substrate (Thermo Scientific), photographed with a ChemiDoc Imaging system, and quantified with NIH ImageJ.

### Behavioral tests

***Elevated zero maze*** The elevated zero maze for anxiety was performed by placing the mouse into an open quadrant of a continuous circular 5.5 cm-wide track elevated 65 cm above the floor and divided into alternating walled and open quadrants in a dimly lit room, for 6 min. Video recording from above in the last 5 min was automatically scored for time spent in each quadrant (Top Scan software suite, Clever System Inc.).

***Locomotor sensitization*** The mice were tested for sensitized responses to cocaine using a two-injection protocol (Valjent et al., 2010). Briefly, mice were placed in an Accuscan open field chamber (20 × 20 cm^2^, 8 beam sensors for each dimension) to monitor locomotor activity. Room light was set up as 30-50 lux. For the first 4 days, mice were placed for 30 min in the activity chamber (habituation phase), received an injection of saline (*i.p.*), then placed back to the chamber for 1 h. Then locomotor sensitization was tested by replacing injection of saline with cocaine (15 mg/kg, *i.p.*) for two continuous days in the same environment.

***Conditioned Place Preference*** Place-conditioningboxes consistedof two chambers with balanced contexts. A partition with an opening separated the two chambers but allowed animals access to either side. This partition was closed off during the pairing days. The walls, floors and partition of chambers were cleaned with CLIDOX-S and dried after each session. The conditional place preference paradigm was performed using a biased strategy as follows: On day 1, mice were allowed to explore both sides for 15 min, and time spent in each side was recorded. Animals were separated into groups with approximately equalbiases for the sides to start from. The side on which an animal spent most time was defined as “preferred side” for this mouse. On day 2, starting from the unpreferred side, animals were allowed to explore both sides for 15 mins, and data were collected to calculate place preference for the preconditioning phase. Beginning on Day 3, the animals were paired for 8 days (30 min per day), with the saline group receiving injections (0.9% sodium chloride) on both sides of the boxes; whereas the drug-paired group received cocaine (10 mg/kg) on the unpreferred side and saline on the preferred side. On day 11, all the animals were given a saline injection, placed on the unpreferred side, and allowed to run freely between the two sides for 15 min, and time spent on each side was recorded (Top Scan software suite, Clever System Inc.).

### Statistics

Statistical analyses were conducted using Sigma Plot 14.0. Student’s t-tests and factorial model analysis of variances (ANOVA) were employed where appropriate. *Post hoc* analyses were performed using Bonferroni Test or Fisher’s LSD. Data were presented as mean ± s.e.m. Differences were considered to be significant when p<0.05.

## RESULTS

### NCS-Rapgef2 dependency of phospho-ERK activation and Egr-1 induction in response to cAMP elevation in NS-1 cells

The immediate early gene Egr-1/Krox-24/Zif268 is induced by cocaine treatment *in vivo*, and implicated in the behavioral actions of cocaine (Valjent et al., 2006c). We have previously shown that Egr-1 and NCS-Rapgef2 are both required for neuritogenesis induced by Gs-coupled GPCRs including the D1 receptor, or elevation of cAMP itself, in neuroendocrine cells (Ravni et al., 2008; Emery et al., 2013; Emery et al., 2017a; Jiang et al., 2017). In order to link cAMP-dependent activation of NCS-Rapgef2 to signaling to ERK resulting in Egr-1 protein induction, we treated NS-1 rat pheochromocytoma cells, and an NS-1 cell line in which Rapgef2 expression was deleted with CRISPR-mediated gene mutation (Jiang et al., 2017), with 100 μM of the cell-permeant cAMP analog 8-CPT-cAMP, and measured levels of phospho-ERK (pERK) and Egr-1. ERK phosphorylation and Egr-1 induction in response to 8-CPT-cAMP were abolished in the absence of NCS-Rapgef2 (Figure 1). This prompted us to examine whether elevation of both pERK and Egr-1 after cocaine administration, in dopaminoceptive neurons *in vivo*, might also be linked by the cAMP-dependent action of NCS-Rapgef2.

**Figure 1.**
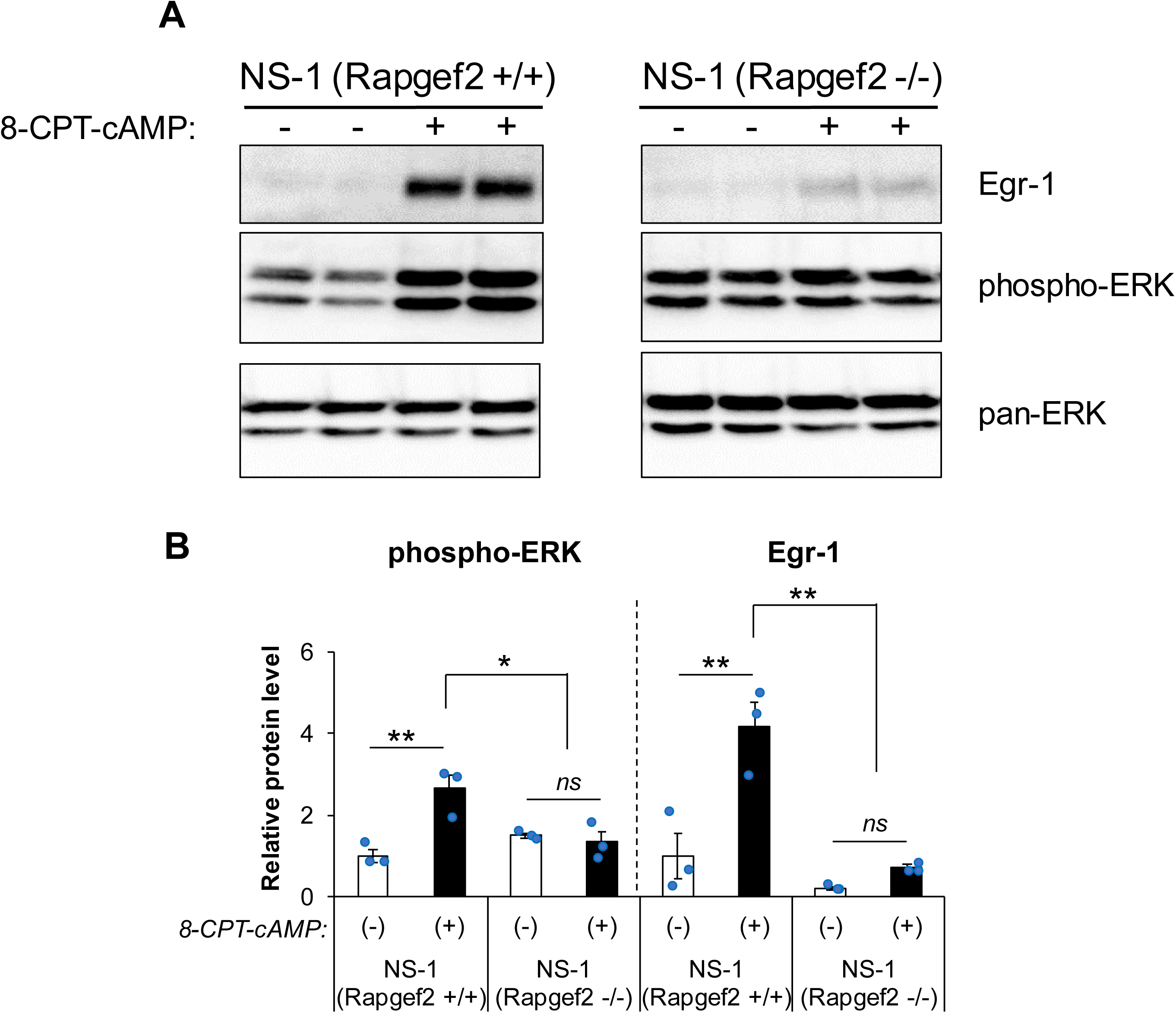
NCS-Rapgef2 dependent phospho-ERK activation and Egr-1 upregulation in neuroendocrine cell line NS-1 induced by 8-CPT-cAMP. **(A)** NS-1 cells (NS-1(Rapgef2 +/+)) or NS-1 cells with NCS-Rapgef2 KO by CRISPR (NS-1 (Rapgef2 -/-)) were treated with 100 µM 8-CPT-cAMP or 0.01% DMSO (8-CPT-cAMP (-) group) for 1 hour, then the cell lysate were collected for western blot analysis of pan-ERK, phospho-ERK and Egr-1. Equal amount of total protein (20 µg) was loaded in each lane. **(B)** Phospho-ERK and Egr-1 protein levels were quantified with Image J and compared to NS1 (Rapgef2+/+)/8-CPT-cAMP (-) group (n=3 per group). *post hoc* Bonferroni test following two-way ANOVA, \**p*<0.05. \*\**p*<0.001.

### Postsynaptic localization of NCS-Rapgef2 in dopaminoceptive regions of mPFC, BLA, and NAc

NCS-Rapgef2 is highly expressed in corticolimbic areas and striatum in mouse brain, including cell types and brain regions (cortical pyramidal cells; hippocampal granule cells of dentate gyrus and pyramidal cells of CA1-3; basolateral amygdalar excitatory neurons; central amygdalar inhibitory neurons; and NAc medium spiny neurons (MSNs) (Jiang et al., 2017)) which receive innervation from dopaminergic neurons of the VTA and substantia nigra. Modulation of neurons in these brain regions by dopamine is implicated in several aspects of learning, memory, stress responding, and reward, and dysfunction of these neurons is similarly implicated in the pathogenesis of a broad range of neuropsychiatric and neurological disorders including Parkinson’s disease, schizophrenia, and drug addiction. We wished to determine if NCS-Rapgef2 was localized within dopaminoceptive neurons at an intracellular location consistent with post-synaptic responses to dopamine receptor activation on cells in these regions. For this purpose, we examined the co- and differential localization of NCS-Rapgef2 and both pre- and post-synaptic markers in these regions. NCS-Rapgef2 immunoreactive (IR) signal is not colocalized with Bassoon, a scaffolding protein of presynaptic active zone, but rather is in close apposition to the Bassoon puncta in mPFC, NAc and BLA (Figure 2A). However, NCS-Rapgef2 IR signal is colocalized with postsynaptic marker PSD95 IR signal in the soma and dendrites of neurons in mPFC pyramidal cells, NAc medium spiny neurons and excitatory neurons in BLA (Figure 2B). These results suggest a post-synaptic location of NCS-Rapgef2 in neurons including those responsive to dopamine.

**Figure 2.**
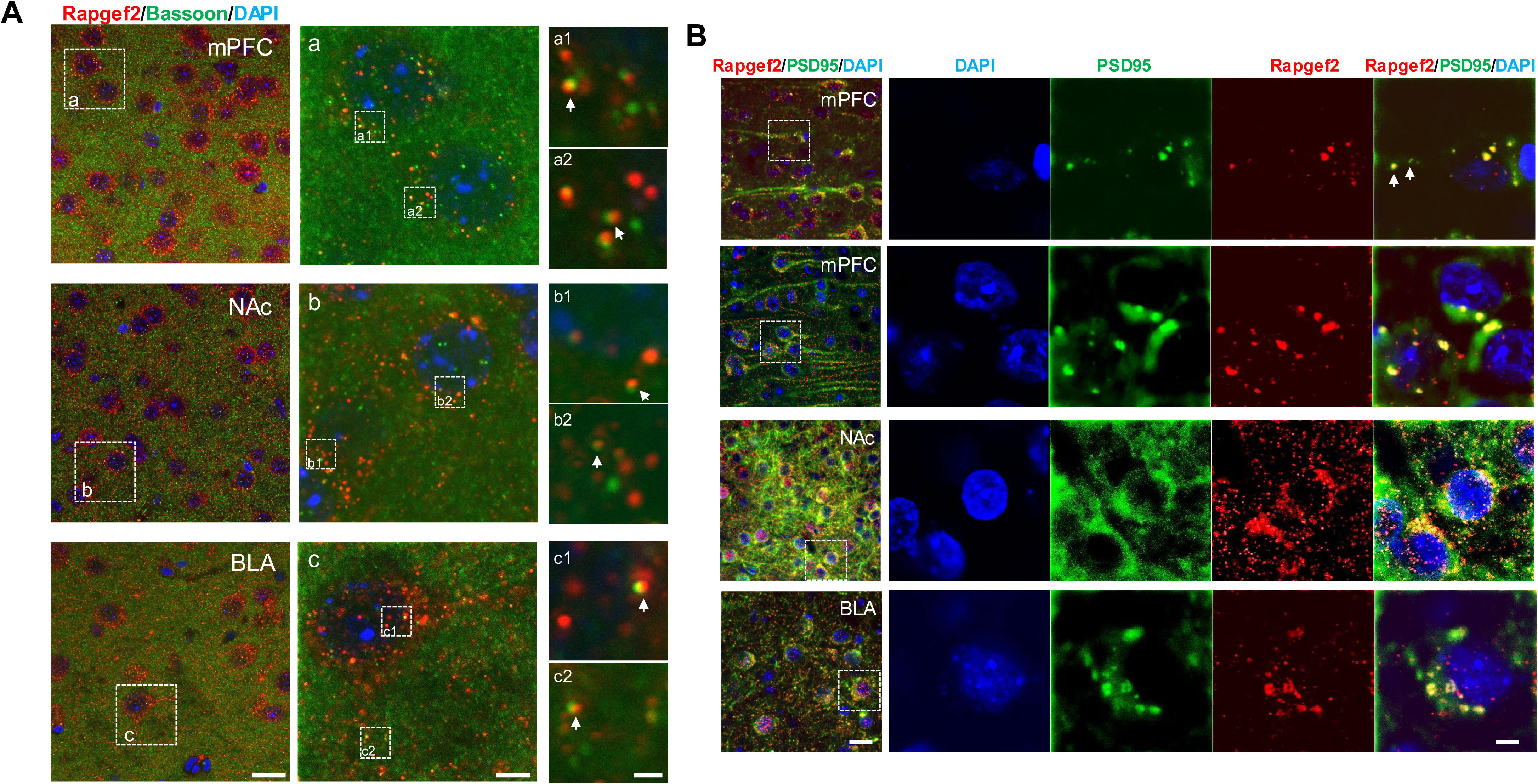
Postsynaptic localization of NCS-Rapgef2 in dopaminoceptive neurons in mPFC, NAc and BLA. **(A)** NCS-Rapgef2 immunoreactive (IR) signal (red) is not colocalized with Bassoon (green), a scaffolding protein of presynaptic active zone, but in close apposition to the Bassoon puncta, indicated by white arrows in a1 & a2 (mPFC), b1 & b2 (NAc) and c1 & c2 (BLA). Scale bar: 20 µm (left); 5 µm (middle); 1 µm (right). **(B)** NCS-Rapgef2 IR signal (red) is colocalized with postsynaptic marker PSD95 IR signal (green) in the soma and dendrites (indicated by white arrows) of neurons in mPFC pyramidal cells, NAc medium spiny neurons and excitatory neurons in BLA. Right panels show, at higher magnification, the images outlined by white boxes on the left. Scale bar: 20 µm (left); 5 µm (right).

### ERK activation by D1 receptor stimulation in mouse nucleus accumbens *ex vivo*: dependence on NCS-Rapgef2

Given the existence of a cyclic AMP→ERK→Egr-1/Zif268 pathway dependent on NCS-Rapgef2 in neuroendocrine cells in culture, we next inquired about the existence of this pathway in neurons receiving dopaminergic input within the mouse CNS, as already implied by our previous observations of Rapgef2-dependent ERK activation in various brain regions after systemic administration of SKF81297 (Jiang et al., 2017). Accordingly, we examined the effects of SKF81297 on pERK activation after its direct application to nucleus accumbens-containing brain slices, in the presence and absence of the NCS-Rapgef2 inhibitor 6-phenyl-9-tetrahydrofuranyl-adenine (EL1101). 10 μM SKF81297 produced a robust increase in pERK in both core and shell of nucleus accumbens slices in a pattern similar to that seen after SKF81297 administration *in vivo* (Figure 3). Furthermore, this activation was completely confined to D1- expressing neurons (i.e. >95% phospho-ERK positive neurons are D1-receptor/tdTomato positive—see Figure 3). SKF81297-stimulated ERK phosphorylation was completely blocked in the presence of 100 μM of EL1101 (Figure 3), indicating that the effects of D1 agonist treatment on ERK activation are mediated directly at D1 dopaminoceptive medium spiny neurons (MSNs) of the NAc, and are confined to these neurons.

**Figure 3.**
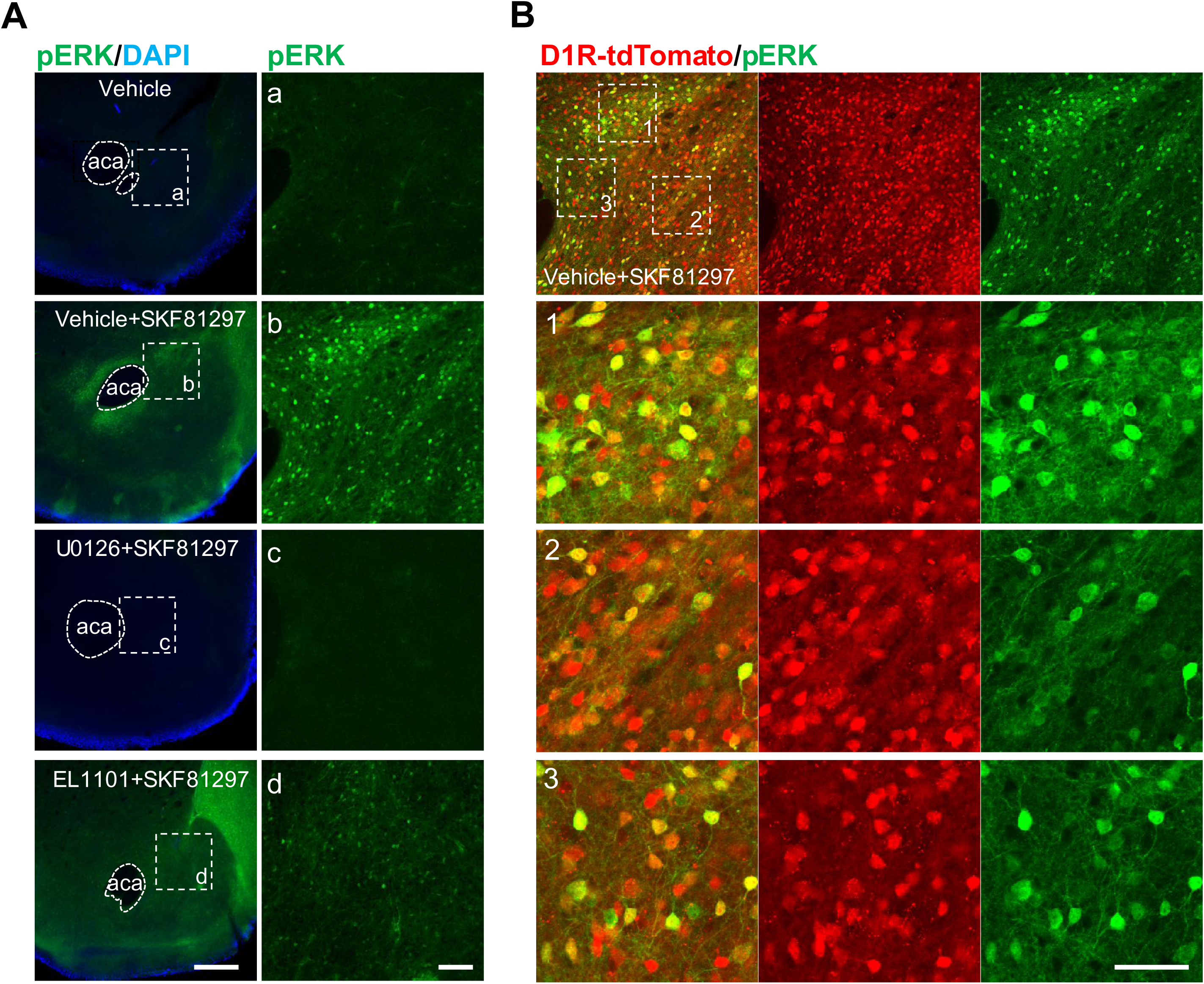
NCS-Rapgef2 inhibitor N6-phenyl, 9-tetrahydrofuranyl adenine (EL1101) attenuated phospho-ERK activation in the NAc of mouse brain slice induced by D1 receptor agonist SKF 81297. **(A)** Brain slices (coronal sections in 250 µm) were incubated vehicle (DMSO final concentration 0.1%), 10 μM SKF81297, 10 μM U0126 plus 10 μM SKF81297 or 100 μ EL1101 plus 10 μ SKF81297 for 30 min. After fixation with 4% PFA, slices were stained with phospho-ERK antibody. Phospho-ERK activationin the NAc of mouse brain slice can be induced by D1 receptor agonist SKF81297, thatwas blocked by the MEK1 inhibitor U0126 or attenuatedby NCS-Rapgef2 inhibitor EL1011. Right panels show, at higher magnification, the images outlined by white boxes on the left. Scale bar: 500 µm (left); 100 µm (right). **(B)** D1-specific activation of phospho-ERK in the NAc of D1d1a-tdTomato mouse indicated by colocalization of phospho-ERK IR signal (green) and D1 MSNs (tdTomato, red). Scale bar: 50 µm (left).

### Cocaine-induced up-regulation of ERK and Egr-1/Zif268 inmouse brain requires expression of NCS-Rapgef2

We next sought to specifically manipulate Rapgef2 levels in dopaminoceptive brain regions, the mPFC, BLA, and NAc, closely associated with behaviors affected by cocaine. NCS-Rapgef2 knock-out specifically in corticolimbic areas, but not nucleus accumbens, was achieved by crossingfloxed Rapgef2 (Rapgef2^cko/cko^) with Camk2α-cre mice. NCS-Rapgef2 expression was decreased at least 85% in mPFC and BLA in these mice, while NCS-Rapgef2 expression in NAc was unaffected(Figure 4A). Knock-down of NCS-Rapgef2 expression in D1 MSNs in NAc is not achievable by crossing a D1 receptor cre driver line with Rapgef2^cko/cko^ (Drd1-cre^+/-^:: Rapgef2^cko/cko^) (Extended Data Figure 4-1) or by crossingother striatum-specific Cre driver lines, such as Gng7-Cre, with Rapgef2^cko/cko^ mice (data notshown). Therefore, an alternative method was employed to achieve the ablation of NCS-Rapgef2 expression in NAc, i.e. by bilateral injection of AAV9-hSyn-Cre-eGFP into the ventral striatum of Rapgef2^cko/cko^ mice (Figure 4B). Expression of NCS-Rapgef2 in the NAc was not affected by injection of AAV9-hSyn-eGFP control virus into Rapgef2^cko/cko^ mice, or by injection of AAV9-hSyn-cre-eGFP into C57BL/6J wild-type mice. However, NCS-Rapgef2 in NAc was significantly reduced by injection of AAV9-hSyn-Cre-eGFP into Rapgef2^cko/cko^ mice (Figure 4C).

**Figure 4.**
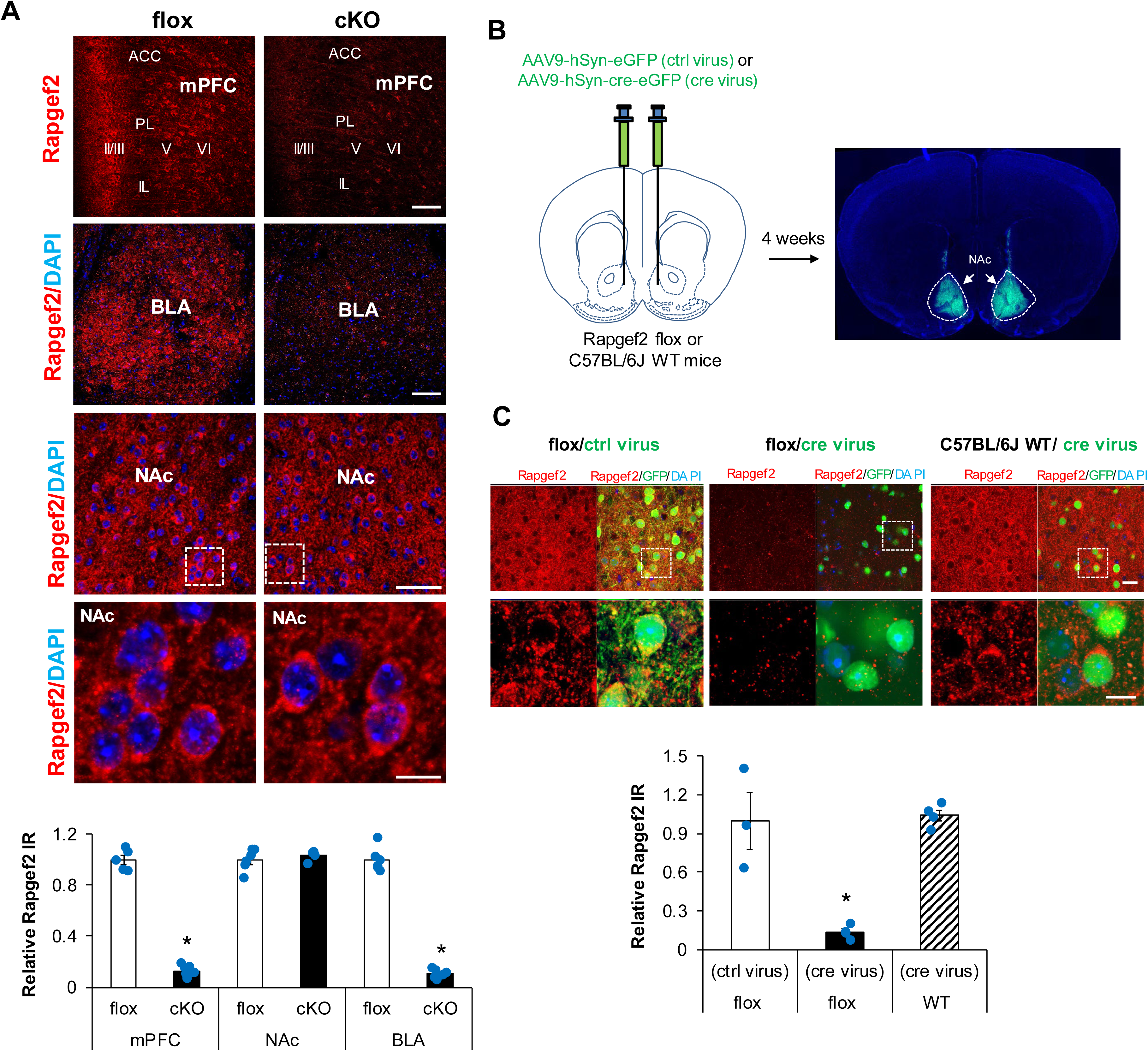
Region-specific ablation of NCS-Rapgef2 expression. **(A)** Camk2α-cre^+/-^::Rapgef2^cko/cko^ (cKO) was generated by crossing Rapgef2^cko/cko^ (flox) with Camk2α-cre mice. NCS-Rapgef2 expression was largely ablated in mPFC and BLA of cKO, while was unaffected in NAc of cKO. Scale bar: 100 µm in mPFC and BLA images; 50 µm & 10 µm in NAc. N=4∼7 animals per group. Student’s t-test for each region, \**p*<0.001. **(B)** Representative image of brain slice 4 weeks after AAV bilateral viral injection in NAc. **(C)** NCS-Rapgef2 in NAc was abrogated by bilateral injection of AAV9-hSyn-cre-eGFP (cre virus) into the ventral striatum of Rapgef2^cko/cko^ (flox) mice. However, expression of NCS-Rapgef2 in the NAc was notaffected by injectionof AAV9-hSyn-eGFP (ctrl virus) into Rapgef2^cko/cko^ (flox) mice or by injection of AAV9-hSyn-cre-eGFP (cre virus) into C57BL/6J wild-type mice. Bottom panels show, at higher magnification, the images outlined by white boxes in upper panels. Scale bar: 20 µm (top) and 10 µm (bottom). NCS-Rapgef2 IR signals in GFP-positive neurons were quantified with ImageJ. N=4∼7 animals per group. *post hoc* Bonferroni test following one-way ANOVA, \**p*<0.05.

Thus, NCS-Rapgef2 expression can be efficiently and specifically eliminated in NAc, by stereotaxic AAC9-hSyn-Cre-eGFP injection in Rapgef2^cko/cko^ mice, and in BLA and mPFC, by genetic cross of Camk2α-cre^+/-^ and Rapgef2^cko/cko^ mice.

Cocaine treatment is known to induce phospho-ERK elevation and trigger ERK-dependent Egr-1/Zif268 expression in D1 medium-sized spiny neurons (Bertran-Gonzalez et al., 2008), implicated in locomotor and drug preference responses to cocaine (Valjent et al., 2006c). To confirm this, and to also confirm the existence of the ERK→Egr-1/Zif268 pathway in D1-dopaminoceptive neurons in mPFC and BLA, we treated mice in which D1 neurons are labeled with tdTomato, with cocaine. Phospho-ERK neuronal induction after cocaine occurs predominantly in D1-dopaminoceptive (tdTomato-positive) neurons in NAc, BLA and mPFC (Figure 5A). We further quantified the percentage of pERK+ cells that were also D1 receptor-expressing (tdTomato-positive) after treatment with cocaine. In prefrontal cortex, 56.7 ± 2.8% of pERK+ neurons in layer II/III, 78.1 ± 2.4% of pERK+ neurons in layer V, and 97.3 ± 0.7% pERK+ neurons in layer VI were also tdTomato-positive. In BLA, 72.2 ± 2.8 % of pERK+ neurons were D1 receptor positive, and in shell and core of NAc almost all (98.6 ± 0.1%) pERK+ neurons were D1-MSNs.

**Figure 5.**
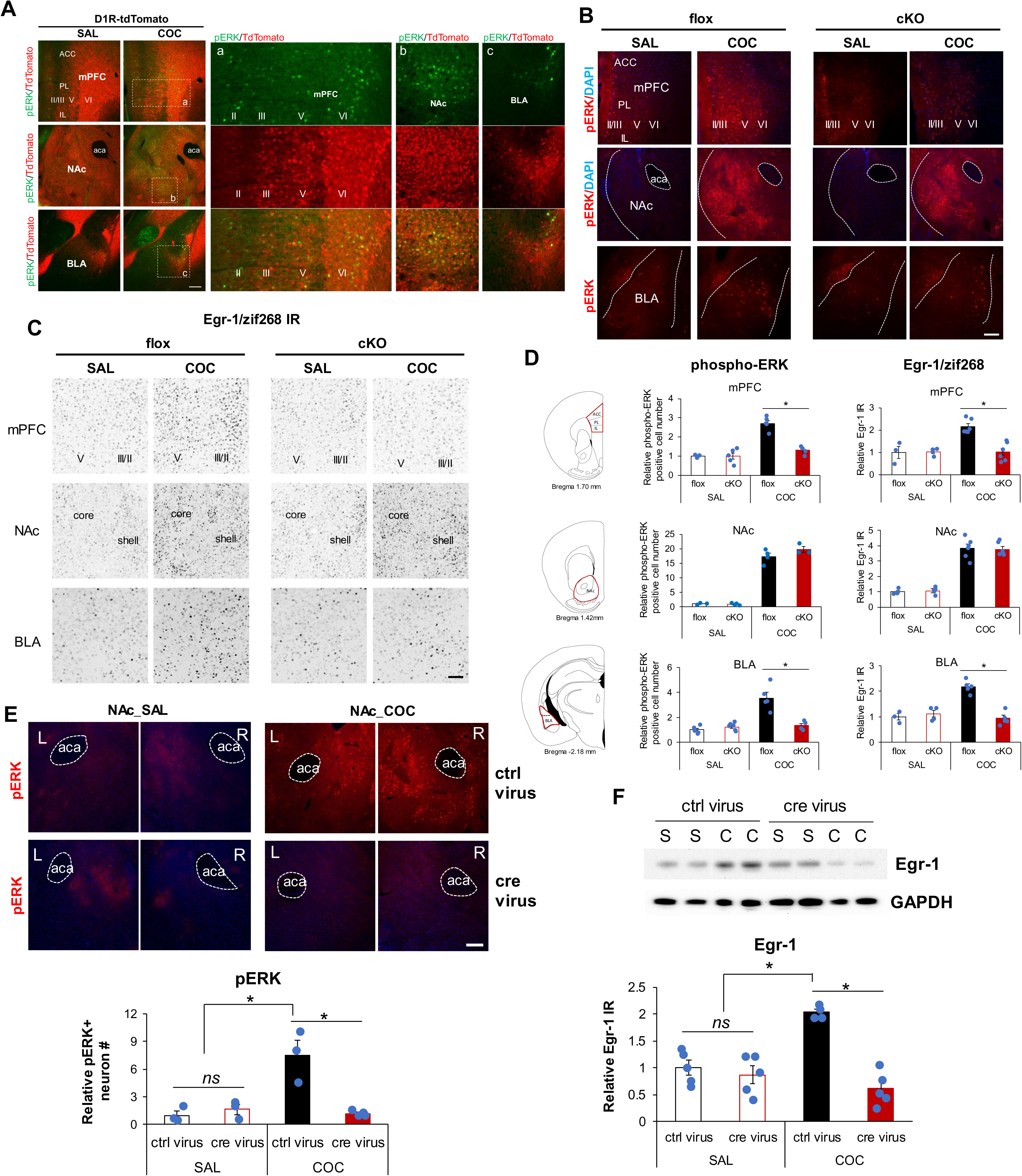
Expression of NCS-Rapgef2 is required for cocaine-induced up-regulation of ERK and Egr1/zif268 in mouse brain **(A)** Majority of cocaine-induced phospho-ERK activation (10 min after administration of 20 mg/kg cocaine, ip) was in D1 receptor positive neurons in mPFC, NAc and BLA. Right panels (indicated as a, b, c) show the images at higher magnification, outlined by white boxes on the left. Scale bar: 200 µm. **(B)** Phospho-ERK elevation observed 10 mins after *ip* administration of 20 mg/kg cocaine in Rapgef2^cko/cko^ (flox) mice, is attenuated in mPFC and BLA, but not in NAc, of Camk2α-cre^+/-^ ::Rapgef2^cko/cko^ (cKO) mice. Scale bar: 200 µm. **(C)** Egr-1/zif268 elevation observed 1 h after 20 mg/kg cocaine treatment in Rapgef2^cko/cko^ (flox) mice is attenuated in mPFC and BLA, but not in NAc, of Camk2α-cre^+/-^::Rapgef2^cko/cko^ (cKO) mice. Scale bar: 100 µm. **(D)** Quantification of phospho-ERK and Egr-1/zif268 in Rapgef2^cko/cko^ (flox) or Camk2α-cre^+/-^::Rapgef2^cko/cko^ (cKO) mice after saline or cocaine treatment. Relative number of phospho-ERK positive neurons or Egr-1 IR from mPFC, NAc and BLA of each animal was normalized by the average value from age-matched saline-treated Rapgef2^cko/cko^ (flox) mice. N=3∼6 animals per group. *post hoc* Bonferroni test following two-way ANOVA, \**p* < 0.05. (phospho-ERK in mPFC: genotype effect F _(1, 13)_ = 21.312, p<0.001, drug effect F _(1, 13)_ = 43.108, p<0.001, interaction F _(1,13)_ = 21.502, p<0.001; phospho-ERK in NAc, genotype effect F _(1, 10)_ = 2.131, p=0.175, drug effect F _(1, 10)_ = 433.739, p<0.001, interaction F _(1, 10)_ = 2.427, p=0.15; phospho-ERK in BLA: genotype effect F_(1, 16)_ = 14.395, p=0.002, drug effect F_(1, 16)_ = 26.668, p<0.001, interaction F_(1, 16)_ = 23.691, p<0.001; Egr-1 in mPFC: genotype effect F _(1, 15)_ = 10.348, p=0.006, drug effect F _(1, 15)_ = 11.435, p=0.004, interaction F _(1, 15)_ = 11.62, p=0.004; Egr-1 in NAc: genotype effect F _(1, 16)_ = 0.00125, p=0.972, drugeffect F_(1, 16)_ = 169.616,p<0.001, interaction F_(1, 16)_ = 0.11, p=0.744; Egr-1 in BLA: genotype effect F _(1, 13)_ = 19.872, p<0.001, drug effect F _(1, 13)_ = 15.786, p=0.002, interaction F _(1,13)_ = 29.799, p<0.001) **(E)** Bilateral ablation of NCS-Rapgef2 in NAc impaired phospho-ERK elevation in response to cocaine. AAV9-hSyn-eGFP (ctrl virus) or AAV9-hSyn-cre-eGFP (cre virus) was injected into the NAc of Rapgef2^cko/cko^ mice. Four weeks later, phospho-ERK activation in both sides of NAc was examined by immunohistochemistry after acute cocaine (*ip*, 20 mg/kg, 10 min) injection. N=3∼4 animals per group. *post hoc* Bonferroni test following two-way ANOVA, \**p* < 0.05. (phospho-ERK in NAc: genotype effect F _(1, 9)_ = 12.262, p=0.007, drug effect F _(1, 9)_ = 13.901, p=0.005, interaction F _(1, 9)_ = 18.216, p=0.002) **(F)** Cocaine-induced Egr-1/zif268 increase in the NAc of Rapgef2^cko/cko^ mice with ctrl virus, was impaired in the NAc with cre virus. Four weeks after viral infusion, mice were sacrificed 1 hr after saline or cocaine (20 mg/kg) injection. The tissue from NAc was punched outfrom 0.5 mm coronal sections of mouse brains and protein lysate was used for western blot with Egr-1/zif268 antibody. Total protein loading of each sample was first normalized by GAPDH immunoreactivity (IR). Relative Egr-1/zif268 IR (means ± SEM) was obtained by comparing to average of that from NAc of saline-treated Rapgef2^cko/cko^ bilaterally injected with AAV9-hSyn-eGFP (ctrl virus). N=4∼5 animals per group. *post hoc* Bonferroni test following two-way ANOVA, \**p* < 0.05. (Egr-1 in NAc: genotype effect F _(1, 15)_ =31.507, p<0.001; drug effect F _(1, 15)_ =7.995, p=0.013; genotype X drug effect F _(1, 15)_ =21.792, p<0.001)

We examined the requirement for NCS-Rapgef2 in cocaine-induced ERK phosphorylation/activation, and subsequent Egr-1/Zif268 induction, in two sets of experiments. First, the effects of cocaine injection were compared in Rapgef2^cko/cko^ control mice, and in Camk2α-cre^+/-^::Rapgef2^cko/cko^ mice in which Rapgef2 expression is eliminated in mPFC and BLA but not in NAc. ERK phosphorylation and Egr-1/Zif268 induction occurred in all three areas in control mice at 10 and 60 min, respectively after *i.p.* cocaine administration (20 mg/kg). Induction of both ERK phosphorylation and Egr-1/Zif268 was abolished in mPFC and BLA, where Rapgef2 expression was absent, butnotin NAc, where Rapgef2 expressionwas spared(Figure 5B-D). Next, the effects of cocaine injection were compared in Rapgef2^cko/cko^ mice receiving AAV-Syn-Cre-eGFP (to eliminate Rapgef2 expression in NAc) or AAV-Syn-eGFP (controls). ERK phosphorylation and Egr-1/Zif268 induction by cocaine were elimninated in NAc after loss of NCS-Rapgef2 expression in mice receiving intra-NAc AAV-Syn-Cre-eGFP injection (Figure 5E-F) with no effect on ERK phosphorylation or Egr-1/Zif268 induction in mPFC or BLA (data not shown). Similar results were obtained after *ip* injection of dopamine D1 receptor agonist SKF81297 (Extended Data Figure 5-1). Thus, expression of NCS-Rapgef2 is required for cocaine-induced up-regulation of ERK and Egr1/Zif268 in D1-expressing neurons of NAc, of mPFC, and of BLA.

We note here that both D1 and D2 MSNs express NCS-Rapgef2 (Extended Data Figure 5-2A). To determine if ERK activation in D2 MSNs requires NCS-Rapgef2, Rapgef2^cko/cko^ mice with NCS-Rapgef2 ablation in the NAc by AAV-Syn-cre-eGFP infusion were treated with the D2 antagonist eticlopride, which caused pERK induction in D2-MSN unaffected by ablation of NCS-Rapgef2 expression (Extended Data Figure 5-2B).

### NCS-Rapgef2 in D1 MSNs of ventral striatum, but not excitatory neurons in mPFC or BLA, contributes to cocaine-induced psychomotor sensitization and conditioned place preference

Neither Camk2α-cre^+/-^::Rapgef2^cko/cko^ mice, nor Rapgef2^cko/cko^ mice infused with AAV-Syn-Cre-eGFP in NAc, showed altered locomotor activity and anxiety in the elevated zero maze, indicated by similar total distance traveled and time in open arms (Extended Data Figure 6-1), compared to each other or to corresponding control mice. Mice were then subjected to testing for locomotor sensitization to cocaine using the two-injection protocol (Valjent et al., 2010)(Figure 6A): For the first 4 days, animals were habituated for 30 mins in an activity box, then injected with saline, and allowed free activity for one hour. On day 5, after 30 min of habituation in the same box in the same room, animals received their first dose of cocaine (15 mg/kg, *ip*). All four groups of animals showed hyperactivity after cocaine injection (Figure 6B and 6C). On day 6, in the same context, animals received a second dose of cocaine (15 mg/kg, *ip*) after the habituation trial. Both control groups (Rapgef2^cko/cko^ mice, and Rapgef2^cko/cko^ mice injected with AAV-Syn-eGFP control virus in the NAc) exhibited a comparable increase in locomotor activity compared with their response to the first dose of cocaine (Figure 6B and 6C). Similar locomotor sensitization in response to cocaine was observed in another control group: C57BL6/J wild-type mice injected with AAV-Syn-cre-eGFP virus in the NAc (Extended Data Figure 6-2). Locomotor sensitization similarly unaffected in Camk2α-cre^+/-^::Rapgef2^cko/cko^ mice. However, cocaine-induced locomotor sensitization was abolished in Rapgef2^cko/cko^ mice with AAV-Syn-cre-eGFP bilateral infusion in the NAc (Figure 6C).

**Figure 6.**
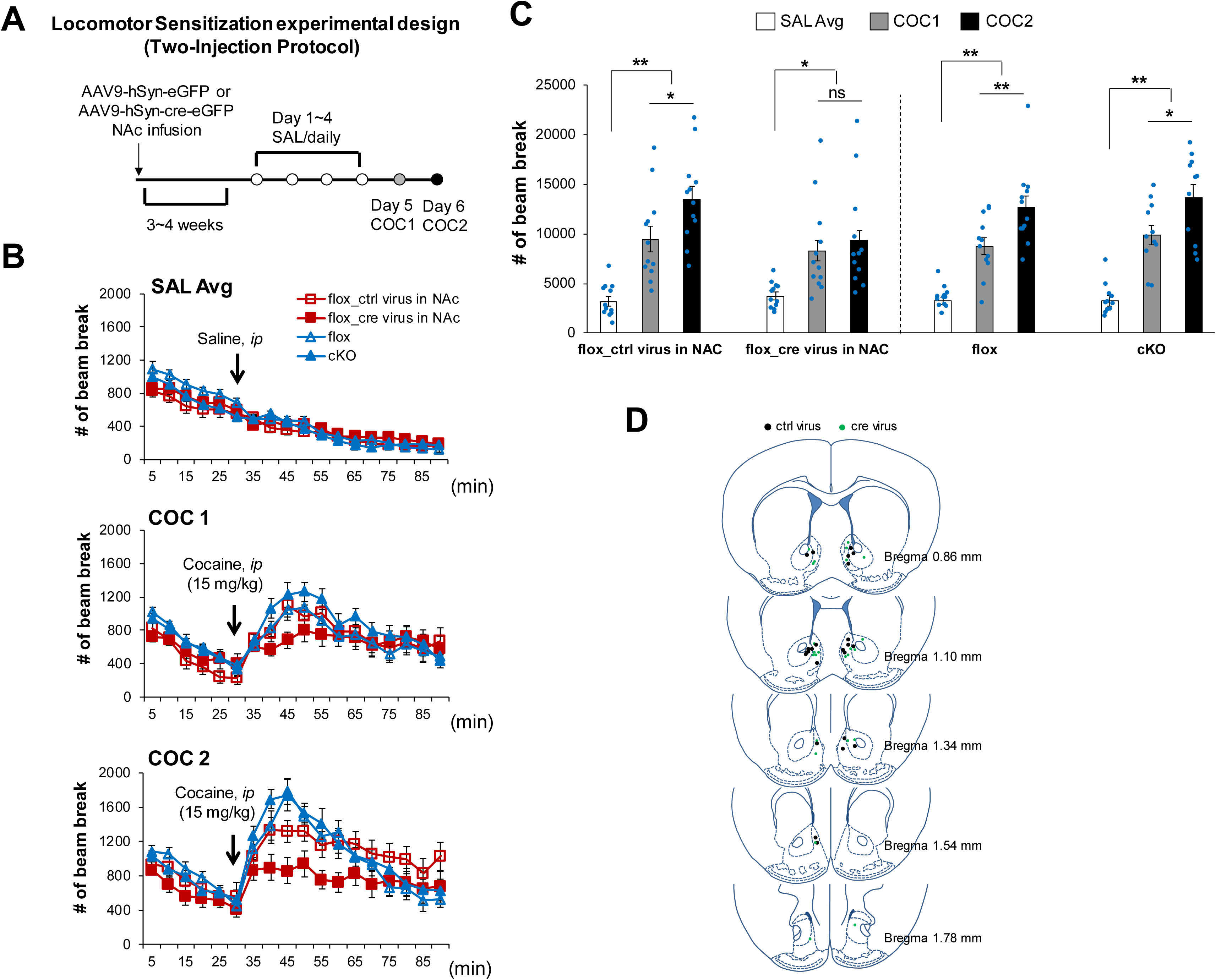
Locomotor sensitization induced by cocaine was abolished in the mice with NCS-Rapgef2 ablation in the NAc. **(A)** Experimental design for locomotor sensitization. Camk2α-cre^+/-^::Rapgef2^cko/cko^ (cKO) mice, Rapgef2^cko/cko^ (flox) mice with bilateral NAc AAV9-hSyn-cre-eGFP (cre virus) injection, and their corresponding controls were subjected to a two-injection protocol (TIPS) for cocaine locomotor sensitization. For the first 4 days, animals were habituated for 30 min in an activity chamber, then injected with saline, and allowed free movement within the chamber for 60 min. On day 5, after 30 min of habituation in the same box in the same room, animals received the first dose of cocaine. On day 6, in the same context(same box in same room), animals receiveda second dose of cocaine after a habituation trial. **(B)** Animal locomotor activity in response to saline or cocaine administration. Activity was monitored in two trials each day: 30 minutes free running and 1 hour after saline or cocaine (15 mg/kg) injection. **(C)** Locomotor activity (total number of beam breaks) for one hour following saline or cocaine administration. SAL Avg, average of activity during 4 days of saline injection; COC1, activity after the first dose of cocaine (15 mg/kg) on day 5; COC2, activity after the second dose of cocaine (15 mg/kg) on day 6. N=12 animals per group. Two-way ANOVA followed by *post hoc* Fisher’s LSD test, *p<0.05 ** p<0.001. (genotype effect: F _(3, 129)_ =1.804, p=0.15; drug effect: F _(2, 129)_ =73.632, p<0.001; interaction: F _(6, 129)_ =1.201, p=0.31. No difference between groups within SAL Avg; No difference between groups within COC1; cre virus group is different from either group within COC2). **(D)** Schematic representation of the injection sites in the NAc for the mice used for locomotor sensitization test. N=12 animals for control virus injection; n=12 animals for cre virus injection.

We also studied the effects of region-specific NCS-Rapgef2 knockout on the effects of repeated administration of cocaine on drug preference, using the conditioned place preference (CPP) test (Figure 7A). Ablation of NCS-Rapgef2 in NAc by viral injection also abolished cocaine-induced conditioned place preference (Figure 7B). These results indicate that two major behavioral consequences of repeated psychomotor stimulant (cocaine) administration are dependenton ERK→Egr-1/Zif268 signaling in D1-MSNs, and further suggestcellular mechanistic linkage between the two behaviors. CPP was not affected in Camk2α-cre^+/-^::Rapgef2^cko/cko^ mice (Figure 7C), indicating that the development of conditioned place preference inheres in cellular changes in dopaminoceptive neurons of the nucleus accumbens, but not those in BLA and mPFC.

**Figure 7.**
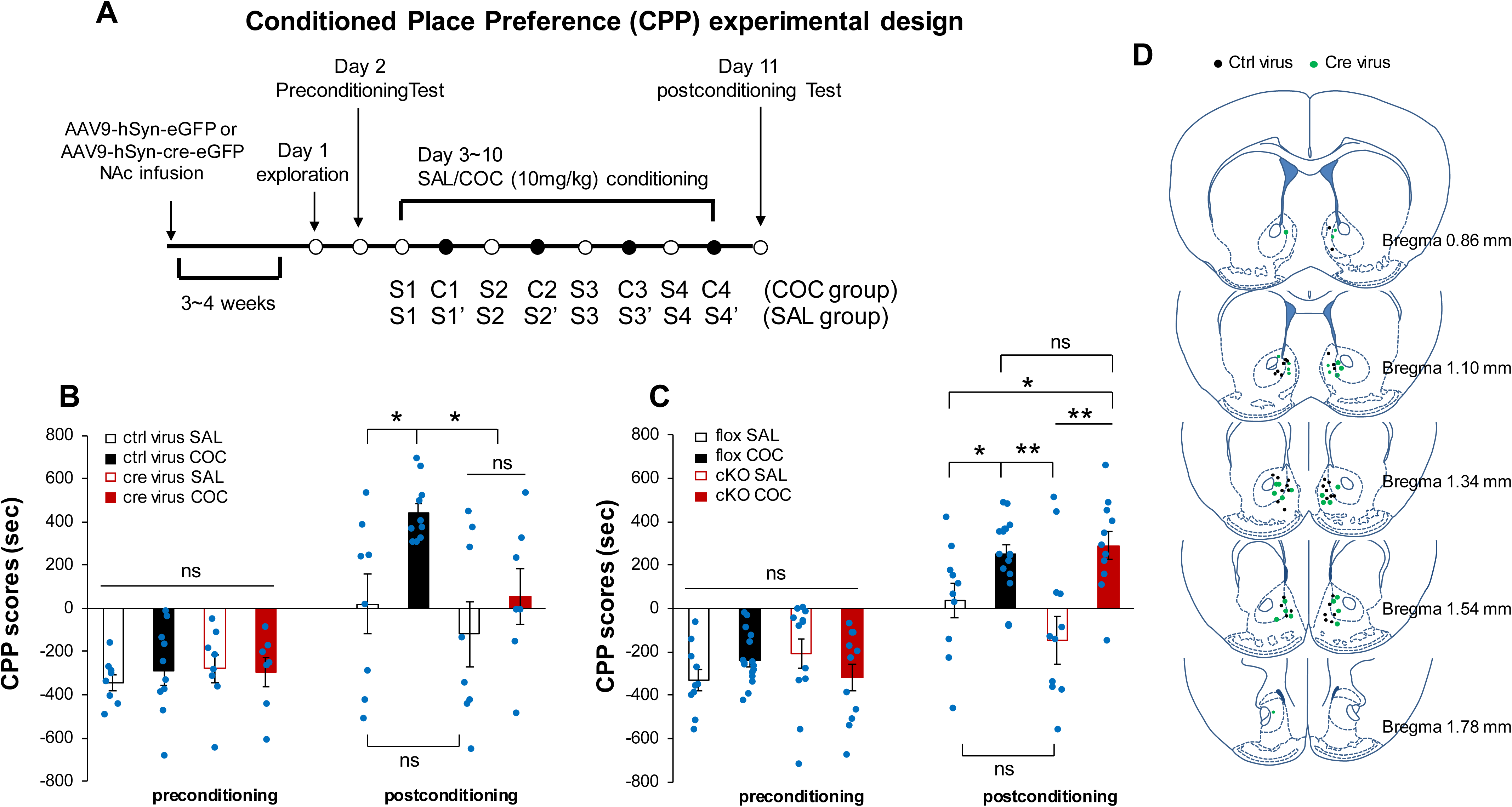
Conditioned place preference (CPP) induced by cocaine was impaired in the mice with NCS-Rapgef2 KO in the NAc. **(A)** Experimentaldesign for CPP. A shuttle box with a door that can be opened to allow the animal free access to the two chambers was used for CPP. The two chambers differed on several sensory and environmental properties. Animals received 8 days /four-sets of cocaine and saline-pairing: for cocaine group, pair one chamber with cocaine, pair opposite one with saline, for saline group, pair both sides with saline. Animals were tested for the place preference before drug pairing and postconditioning by recording the amount of time they spent in each chamber. **(B)** CPP scores for mice with NCS-Rapgef2 KO in the NAc compared to the corresponding controls. N=7∼10 animals per group. *post hoc* Bonferroni test following three-way ANOVA, *p<0.05. (pre- vs post-conditioning: F _(1, 58)_ =36.937, p<0.001; genotype effect: F _(1, 58)_ =3.059, p=0.086; drug effect: F _(1, 58)_ =5.687, p=0.02; conditioning X genotype interaction: F _(1, 58)_ =4.875, p=0.031; conditioning X drug interaction: F _(1, 58)_ =4.5, p=0.038; conditioning X genotype X drug interaction: F _(1, 58)_ =0.443, p=0.508). **(C)** CPP scores for Camk2α-cre^+/-^::Rapgef2^cko/cko^ (cKO) mice compared to the corresponding controls Rapgef2^cko/cko^ (flox). N=10∼16 animals per group. *Post hoc* Bonferroni test following three-way ANOVA, *p<0.05. **p<0.001. (pre- vs post-conditioning: F _(1, 90)_ =67.764, p<0.001; genotype effect: F _(1, 90)_ =0.315, p=0.576; drug effect: F _(1, 90)_ =11.629, p<0.001; conditioning X genotype interaction: F _(1, 90)_ =0.969, p=0.328; conditioning X drug interaction: F _(1, 90)_ =12.82, p<0.001; genotype X drug interaction: F _(1, 90)_ =0.0099, p=0.921; conditioning X genotype X drug interaction: F _(1, 90)_ =5.332, p=0.023) **(D)** Schematic representation of the injection sites in the NAc for the mice used for CPP test. N=18 animals for ctrl virus injection: n=8 animals for saline group, n=10 animals for cocaine group. N=15 animals for cre virus injection: n=8 animals for saline group, n=7 animals for cocaine group.

### Parcellation of cAMP signaling to ERK and CREB in nucleus accumbens

Several models for intracellular signaling underlying dopamine-dependent ERK activation in D1 dopaminoceptive neurons emphasize that the sole cAMP effector for this process is PKA, both because inhibition of ambient ERK phosphatase activity can be achieved through PKA-dependent D1 signaling, and because previously no direct pathway for ERK activation by cAMP elevation has been identified in neurons. To test whether or not the activation of ERK associated with upregulation of Egr-1/Zif268 and locomotor sensitization/place preference conditioning by cocaine involved activation of both PKA and NCS-Rapgef2, we quantified phospho-ERK and phospho-CREB, as surrogates for NCS-Rapgef2 and PKA activation respectively, after cocaine treatment in Rapgef2^cko/cko^ control and NCS-Rapgef2-deficient mice. As shown in Figure 8, NCS-Rapgef2 knockout in the NAc caused loss of ERK phosphorylation after cocaine treatment, while cocaine-dependent CREB phosphorylation was preserved. Thus, PKA signaling is apparently intact in mice in which both D1-MSN ERK activation, and acquisition of cocaine locomotor sensitization and conditioned place preference, are lost after deletion of NCS-Rapgef2 in NAc.

**Figure 8.**
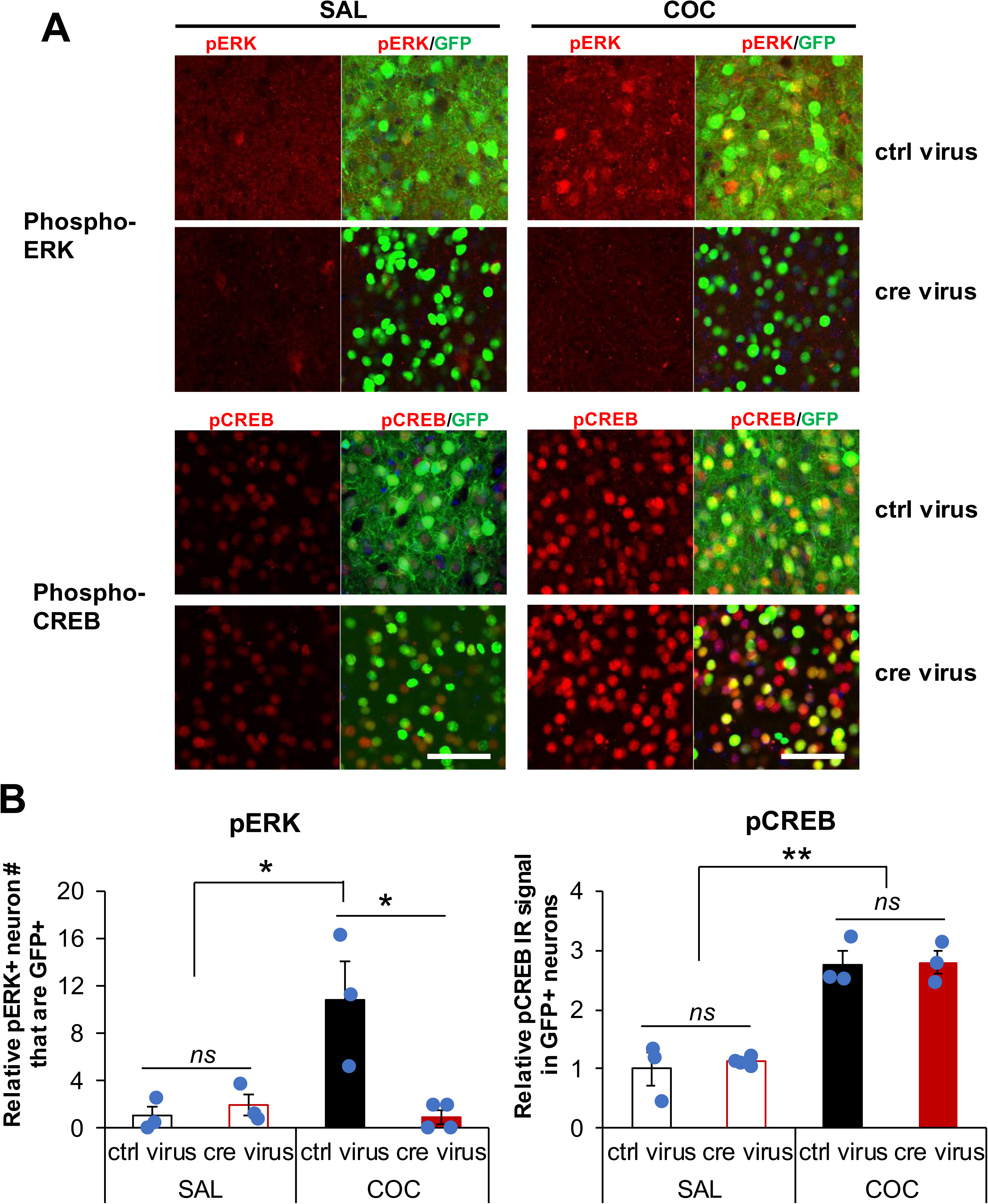
Cocaine treatment resulted in ERK phosphorylation in an NCS-Rapgef2 dependent manner, while CREB phosphorylation was unaffected by NCS-Rapgef2 ablation. AAV9-hSyn-eGFP (ctrl virus) or AAV9-hSyn-cre-eGFP (cre virus) was injected into both sides of NAc of Rapgef2^cko/cko^ mice. After four weeks, saline or cocaine (20 mg/kg) was injected (*ip*) and animals were perfused 10 min later for phospho-ERK and phospho-CREB immunohistochemistry. The number of phospho-ERK/GPF co-positive neurons was significantly increased in the NAc with ctrl virus after cocaine treatment, but not in the NAc with cre virus (n= 3∼4 animals per group; genotype effect F_(1, 9)_ = 7.84, p=0.021; drug effect F_(1, 9)_=7.716, p=0.021; genotype X drug effect F _(1, 9)_ =11.374, p=0.008, two-way ANOVA). Phospho-CREB elevation induced by cocaine treatment was not affected by NCS-Rapgef2 ablation in the NAc (n= 3∼4 animals per group; genotype effect F_(1, 9)_ = 0.173, p=0.687; drug effect F_(1, 9)_=84.582, p<0.001; genotypeX drugeffect F _(1, 9)_ =0.058, p=0.815, two-way ANOVA). Two-way ANOVA followed by *post hoc* Bonferroni test, \**p* < 0.05, **<0.001. Scale bars: 50 µm.

## DISCUSSION

Activation of ERK *in vivo* by D1 agonists. or the psychomotor stimulants cocaine and amphetamine, in D1 neurons is well established (Valjent etal., 2000; Gutierrez-Arenas et al., 2014; Jin et al., 2019). Expression of locomotor sensitization and CPP after cocaine administration is ERK-dependent (Brami-Cherrier et al., 2005; Miller and Marshall, 2005; Ferguson et al., 2006; Valjent et al., 2006b; Valjent et al., 2006a; Fasano et al., 2009), and is linked to ERK-dependent transcription of the *egr1* gene and expression of Egr1/Zif268 protein in dopaminoceptive neurons in striatum and cerebral cortex (Valjent et al., 2006c; Gangarossa et al., 2011).

Whether dopamine signaling to ERK in D1 neurons is direct, or indirect via modulation of ERK activation by other neurotransmitters including glutamate (Valjent et al., 2005) has been an unresolved question. It has been hypothesized that glutamatergic activation of a ras-signaling pathway for ERK phosphorylation is augmented by D1-dependent inhibition of ERK dephosphorylation, shifting the balance between ERK activation/phosphorylation and dephosphorylation towards phospho-ERK elevation and stimulation of ERK-dependent cellular activation events, including *egr1* gene transcription (Valjent et al., 2005; Lu et al., 2006; Santini et al., 2007; Zhang et al., 2010; Santini et al., 2012). This mechanism has been favored because striatal dopaminoceptive neurons possess a PKA- and dopamine responsive protein phosphatase (DARPP-32) which disinhibits the striatally-enriched phosphatase STEP, leading to inhibition of the putative proximate ERK phosphatase PP1 (Valjent et al., 2005; Sun et al., 2007; Gutierrez- Arenas et al., 2014). However, differences between D1 signaling to ERK in ventral and dorsal striatum (Gerfen et al., 2008) have occasioned a search for a direct intracellular signaling pathway for ERK phosphorylation upon dopamine-stimulated elevation of intracellular cAMP. Indeed, G protein-mediated cAMP pathway upregulation has been hypothesized as a common mechanism of adaptation to chronic administration of drugs of abuse (Nestler, 2016), and ERK activation in dopamine D1 receptor-expressing neurons in mesocorticolimbic projection areas, including both medial prefrontal cortex and nucleus accumbens has been with drug-induced neuroadaptations (Valjent et al., 2000; Adams and Sweatt, 2002; Valjent et al., 2005).

We have recently identified intracellular cAMP-dependent signaling downstream of Gs- coupled receptors in neurons and endocrine cells via parallel and insulated (parcellated) pathways initiated by the cAMP effectors PKA, Epac2, and NCS-Rapgef2. These effectors activate distinct but coordinated cellular tasks. Thus, in neuroendocrine NS-1 cells, PKA signaling to CREB mediates cell survival and neuron-specific gene expression; Epac2 signaling to the MAP kinase p38 mediates growth arrest; and NCS-Rapgef2 signaling to the MAP kinase ERK mediates neuritogenesis (Emery et al., 2013; Emery et al., 2014; Emery et al., 2017a). As the ERK pathway is associated with neuritogenesis in NS-1 cells, and similar morphological changes occur via D1 signaling in NAc after cocaine (Lee et al., 2006), we hypothesized that the cAMP→NCS-Rapgef2→ERK pathway might link D1 receptor activation in response to cocaine, to ERK activation and downstream events involved in cellular plasticity causing behavioral effects of repeated psychomotor stimulant administration.

The demonstration that NCS-Rapgef2 is required for Egr-1/Zif268 induction by cyclic AMP in neuroendocrine cells, coupled with our previous finding of NCS-Rapgef2 dependence for dopamine/D1 activation of neuritogenesis in these cells, suggests the existence of a direct dopamine→D1→cAMP→NCS-Rapgef2→ERK→Egr1/Zif268 signaling pathway with potential relevance to cellular signaling in D1-MSNs *in vivo*. A role for NCS-Rapgef2 in mediating the effects of increased synaptic dopamine after cocaine administration is further supported by the localization of NCS-Rapgef2 to post-synaptic elements at dopaminergic synapses, and by NCS-Rapgef2-dependent activation of ERK by the D1 agonist SKF81297 in NAc brain slices *ex vivo*, as reported here. Finally, we have studied the behavioral effects of cocaine in mice in which NCS-Rapgef2 expression was eliminated in various brain regions (mPFC, BLA and NAc) in which dopamine acts at D1 receptors. Loss of NCS-Rapgef2 expression in mPFC and BLA excitatory neurons eliminated ERK activation by both SKF81297 and cocaine in these brain regions, without affecting cocaine-induced locomotor sensitization, or conditioned place preference. Elimination of NCS-Rapgef2 expression in NAc correspondingly abolished ERK activation by cocaine or D1 agonist, and also cocaine-induced locomotor sensitization and conditioned place preference. This occurred without affecting cocaine-induced activation of CREB phosphorylation in MSNs. It remains to be tested whether or not NCS-Rapgef2-dependent signaling in either mPFC or BLA is required for cocaine-dependent behaviors not studied here, such as relapse (Rebec and Sun, 2005).

These findings have several implications for understanding the role of cyclic AMP in mediating the post-synaptic actions of dopamine and subsequent dopamine-dependent behaviors, and for definingthe brain circuits in which dopamine signaling is critical for the various behavioral consequences of acute and chronic cocaine ingestion.

Cocaine reinstatement depends on different cAMP-dependent pathways as a function of the reinstatement stimulus: it is CREB-dependent for cocaine itself, and ERK-dependent for restraint stress (Kreibich and Blendy, 2004; Briand and Blendy, 2013). These findings accord well with our observation of parcellation of cAMP signaling to ERK and CREB through NCS-Rapgef2 and PKA, respectively, in cultured neuroendocrine cell lines (Emery et al., 2014). The present reportdemonstrates thatparcellation extends to D1-dependentactivation of Egr-1/Zif268 via ERK independently of CREB in NS-1 cells. Dopamine-dependent activation of ERK is likewise NCS-Rapgef2-dependent, while signaling to CREB is not. This observation has allowed us to determine that activation of ERK, apparently independently of PKA/CREB, and in dopaminoceptive neurons of the NAc rather than mPFC or BLA, is required for the behavioral effects of repeated cocaine administration.

ERK signaling itself is required for the sensitized locomotor, but not the initial locomotor effects of cocaine administration (Valjent et al., 2006b). NCS-Rapgef2, however, may be an even more practical target for mitigation of the neuroplastic effects of cocaine that underlie its addictive potential, since it represents a single conditional input to ERK activation (see model, Figure 9). A related member of the protein family presentin brain, Rapgef6, mediates a differentsetof memory-related functions in amygdala and hippocampus (Levy et al., 2015) and, interestingly, both Rapgef2 and Rapgef6 are genetically linked to susceptibility to schizophrenia (Maeta et al., 2018). The specialized behavioral functions of Rapgefs in the central nervous system, and the specific pharmacological and behavioral effects of NCS-Rapgef2 demonstrated here, suggests that NCS-Rapgef2 merits consideration as a target for other reward-related psychopathology besides addiction.

**Figure 9.**
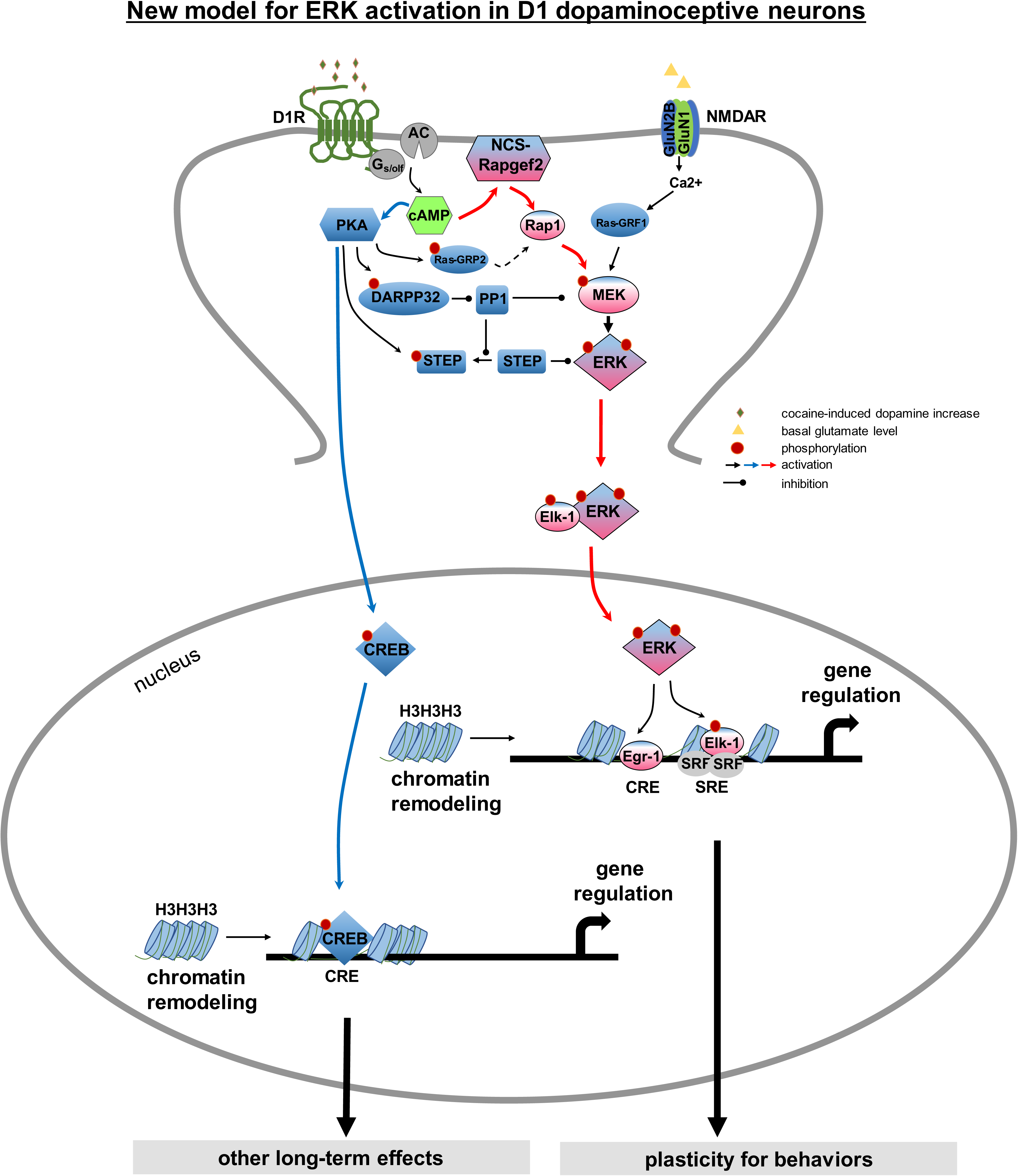
**Proposed direct model for dopamine-dependent ERK activation in D1-dopaminoceptive neurons.** Model for convergent ERK activation by D1 receptor activation adapted from summary of previous conceptualization of this signaling network by Pascoli et al. (Pascoli et al., 2014). Psychomotor stimulants such as cocaine act by increasing synaptic dopamine leading to ERK activation in nucleus accumbens which is required for long-term behavioral adaptations such as locomotor sensitization and CPP. It has previously been thought that dopamine acting through the D1 receptor affects ERK activity only *indirectly,* via PKA- and DARPP-32/STEP-mediated inhibition of ERK dephosphorylation (Svenningsson etal., 2004; Valjent etal., 2005), while *direct* ERK activation itself occurs in the D1 medium spiny neuron only via NMDA receptor-dependent glutamatergic signaling (Pascoli et al., 2011; Cahill et al., 2014). Calcium influx via NMDARs activates calcium-sensitive Ras-guanine nucleotide releasing factor (Ras-GRF1) that in turn can activate the Ras-Raf-MEK cascade (Farnsworth et al., 1995; Fasano et al., 2009). According to this model, the summation of ERK activation by glutamate and of phospho-ERK stabilization by dopamine activity leads to activity-dependent ERK translocation to the nucleus, where it controls both chromatin remodeling and gene transcription, with one of its targets being MSK1/CREB. This model for dopamine signaling in D1-MSNs requires revision (see discussion). First Nagai and colleagues have demonstrated that dopamine can stimulate Rap1 GEF (Rasgrp2) phosphorylation via PKA in MSNs of ventral striatum, implying that ERK could be activated directly by dopamine through Rap1 (Nagai et al., 2016b; Nagai et al., 2016a). We propose here a more parsimonious mechanism, and one more directly supported by experimental evidence, for direct activation of ERK by D1-dependent dopamine signaling, via cAMP activation of NCS-Rapgef2. Thus, we demonstrate here that NCS-Rapgef2 elimination from MSNs in NAc abolishes dopamine-D1-cAMP-dependent activation of ERK, while dopamine-D1-cAMP-PKA-dependent activation of CREB proceeds intact. Thus, cAMP elevation by dopamine in D1-MSNs results not in activation of ERK and CREB in series, but as a *parcellation* of signaling between ERK and CREB resulting in separate cellular consequences under the control of each pathway. We note that our finding of a direct pathway for D1 activation of ERK through NCS-Rapgef2 does not contradict evidence for an indirect modulatory role for PKA-dependent ERK phosphatase inhibition after psychomotor stimulant administration (Valjent et al., 2005; Sun et al., 2007), and is also not inconsistent with additional ERK regulation by glutamatergic input to D1 dopaminoceptiveneurons in the contextof cellular plasticity underlyingcocaineaddiction (Valjent et al., 2000; Park et al., 2013; Cahill et al., 2014; Pascoli et al., 2014). However, it may explain the relative mild, yet more pleiotropic effect(s) of DARPP-32 knock-out (Svenningsson et al., 2004), while the effects of NCS-Rapgef2 are both more marked, and more specific. We posit that D1 receptor activation, and cAMP elevation, in D1 MSNs likely results in parallel effects on ERK, both directly via NCS-Rapgef2 and indirectly via PKA. In this model, the effects of cocaine rely on multiple necessary, but perhaps individually insufficient inputs activated by dopamine, that converge on D1-MSN ERK phosphorylation. These inputs include dopamine action through cyclic AMP and PKA/DARPP/STEP/PP1 (Svenningsson et al., 2004), through cyclic AMP and PKA/RasGRP2/Rap1 (Nagaiet al., 2016b), a src-dependent non-cAMP-mediated augmentation of NMDA receptor-dependent ERK activation via ras (Fasano et al., 2009; Pascoli et al., 2011), and a direct pathway to ERK activation by D1 receptor activation via cyclic AMP and NCS-Rapgef2/Rap1/B-Raf/MEK.

Convergent activation of ERK activation in D1-MSNs through direct (NCS-Rapgef2-mediated) and indirect (PKA-mediated) dopaminergic signaling, and via glutamatergic signaling, provides a mechanistically complete picture of D1-MSN cellular plasticity during repeated exposure to psychomotor stimulants. It has implications for understanding, and intervening in, psychomotor stimulant addiction. In particular, the model (Figure 9) implies that while blockade of D1 signaling altogether could lead to abolition of cocaine-induced locomotor activation, as occurs in D1 receptor deficient (Xu et al., 1994) or D1-MSN ERK-deleted mice (Brami-Cherrier et al., 2005; Ferguson et al., 2006), inhibition of *either* the NCS-Rapgef2 or PKA parcellated signaling pathways may lead to selective blockade of different subsets of learned behaviors occurring during chronic cocaine administration.

Cocaine administration causes ERK activation predominantly if not exclusively in D1-compared to D2-MSNs in the ventral striatum (Valjent et al., 2005; Bertran-Gonzalez et al., 2008), and D1 receptor-expressing neurons play a predominant role in cocaine sensitization and reward learning (see (Baik, 2013) and references therein). Recent studies using optogenetic manipulation and *in vivo* calcium imaging have confirmed the view that activation of D1 neurons promotes the formation of cocaine reward (Lobo et al., 2010; Kravitz et al., 2012; Chandra et al., 2013; Calipari et al., 2016). NCS-Rapgef2 is expressed in both D1 and D2 neurons (Extended Data Figure 5-1). While ERK activation does not occur in D2-MSNs after cocaine administration, it is activated in in this neuronal population by the D2 receptor antagonist eticlopride. However, this response was notimpaired in NCS-Rapgef2-deficient mice (Extended Data figure 5-2). Thus, the NCS-Rapgef2-dependent activation of ERK in D1-MSN signaling documented here is likely to be the major action of this signaling molecule in mediating the cocaine-dependent behaviors examined here. D1- and D2-MSNs in NAc can drive both reward and aversion, depending on their stimulation pattern (Soares-Cunha et al., 2019): it remains to be determined whether or not NCS-Rapgef2 signaling in D2 neurons contributes to these other aspects of NAc function.

Additional questions remain: what is the role of the NCS-Rapgef2 pathway in D1 signaling outside of the NAc? Might signaling through NCS-Rapgef2 mediate cocaine relapse associated with dopamine neurotransmission in mPFC (Miller and Marshall, 2005; Lu et al., 2006; Girault et al., 2007)? What might be the role(s) of CREB-dependent signaling (Shaywitz and Greenberg, 1999), spared upon abrogation of NCS-Rapgef2, in the D1-MSN response to cocaine? Finally, might it be possible to develop brain-permeant NCS-Rapgef2 antagonists of sufficientpotencyand selectivity for use *in vivo*? If dendritic spinogenesis in D1-MSNs is the morphological correlate of Rapgef2-dependent ERK signaling to Egr-1/Zif268 resulting in neuritogenesis in NS-1 cells, it might be supposed that blocking this process could affect not only short-term effects of repeated cocaine administration including locomotor sensitization and cocaine preference, but also longer-term effects of cocaine-seeking behavior associated with altered D1-MSN spine density (Lee et al., 2006; Ren et al., 2010).

## CONFLICT OF INTEREST

The authors declare nocompetingfinancialinterests.

## ACKNOWLEDGEMENTS

Thanks to Haiying Zhangfor mouse colony management andgenotyping. Thanks to Dr. Kazu Nakazawa (The Southern Research Institute) for helpfulcomments. This work was supportedby the National Institute of Mental Health Intramural Research Program, Project 1ZIAMH002386 to L.E.E.

---

**Extended Data Figure 4-1.**
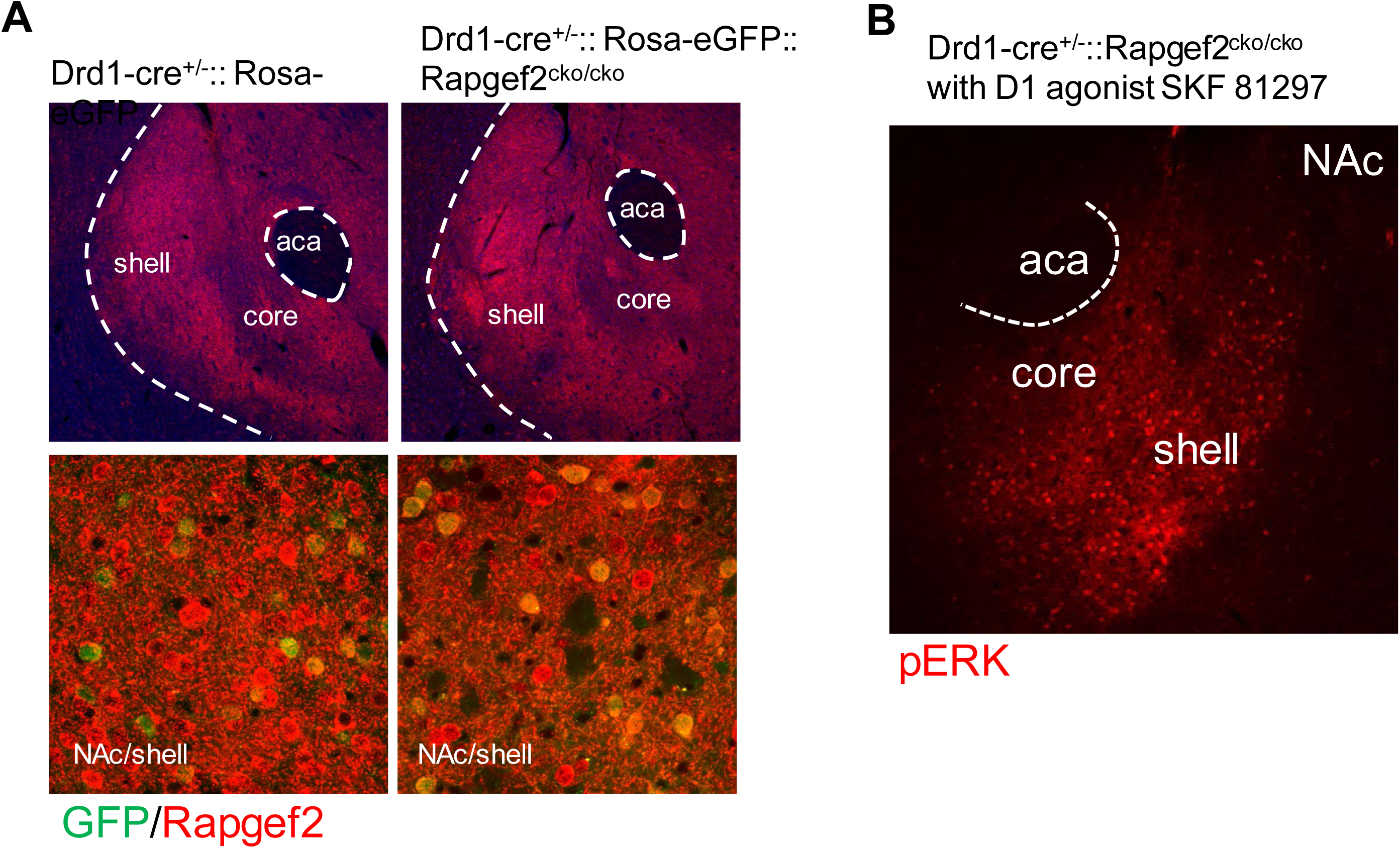
Cre expression in D1d1a-cre mouse line is not strong enough to knockout NCS-Rapgef2 in D1 MSNs in NAc. **(A)** NCS-Rapgef2 expression is largely intact in the NAc (top), especially in D1-MSNs (bottom, in green) of Drd1-cre^+/-^:: Rosa-eGFP:: Rapgef2^cko/cko^ compared to Drd1-cre^+/-^::Rosa-eGFP, indicated by Rapgef2 antibody staining. **(B)** Phospho-ERK activation in the NAc of Drd1-cre^+/-^:: Rapgef2^cko/cko^ was normally induced by D1 receptor agonist SKF81297 (*ip*, 2 mg/kg, 15 min).

**Extended Data Figure 5-1.**
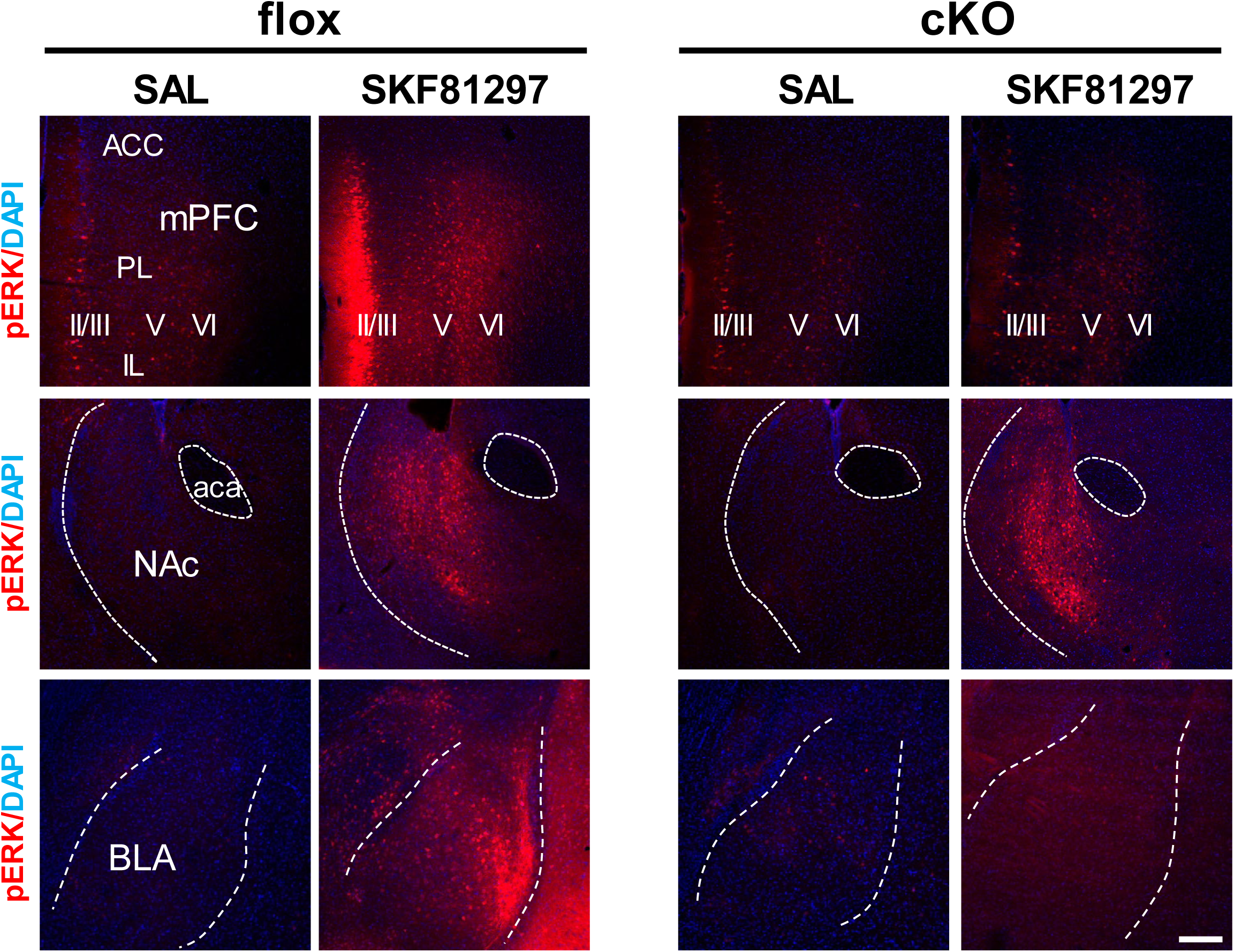
Phospho-ERK elevation observed 30 mins after *ip* administration of 5 mg/kg SKF81927 in Rapgef2^cko/cko^ (flox) mice, is attenuated in mPFC and BLA, but not in NAc, of Camk2α-cre^+/-^::Rapgef2^cko/cko^ (cKO) mice. Scale bar: 200 µm.

**Extended Data Figure 5-2.**
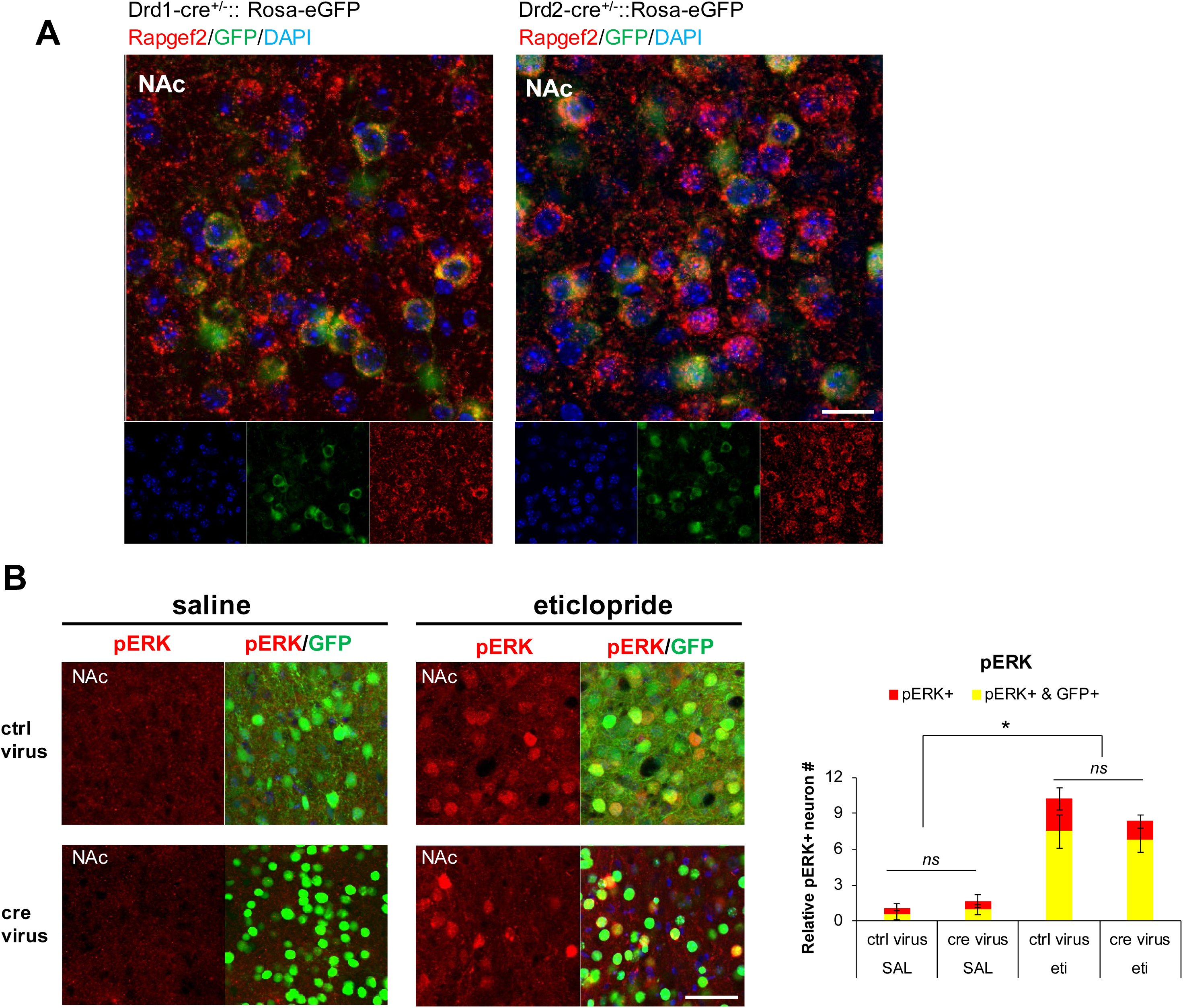
**(A**) NCS-Rapgef2 is expressed in both D1 and D2 MSNs in NAc. NCS-Rapgef2 immunohistochemistry was performed usingbrain sections from Drd1-cre^+/-^::Rosa-eGFP or Drd2-cre^+/-^::Rosa-eGFP mice. NCS-Rapgef2 IR signal (red) was seen in either D1 MSNs, indicated by GFP IR signal (green) in the NAc of Drd1-cre^+/-^::Rosa-eGFP mice, or in D2 MSNs indicated by GFP IR signal (green) in the NAc of Drd2-cre^+/-^::Rosa-eGFP mice. **(B)** pERK induction in presumptive D2-MSNs in NAc after *in vivo* eticlopride administration is not NCS-Rapgef2-dependent. Phospho-ERK activationin NAc was measured in NAc of Rapgef2^cko/cko^ mice in which either AAV9-hSyn-eGFP (ctrl virus) or AAV9-hSyn-cre-eGFP (cre virus) was injected into NAc, and four weeks later mice were treated with either saline or the D2R antagonist eticlopride (2 mg/kg, ip, 15 min). Total number of phospho-ERK positive neurons and phospho-ERK positive neurons with GFP positive (pERK+ & GFP+) in NAc were quantified with ImageJ. Relative number of phospho-ERK positive neurons in NAc of each animal was normalized by the average value from saline-treated Rapgef2^cko/cko^ (flox) mice with ctrl virus. N=3 animals per group. *post hoc* Bonferroni test following two-way ANOVA, \**p* < 0.05. Scale bars: 50 µm.

**Extended Data Figure 6-1.**
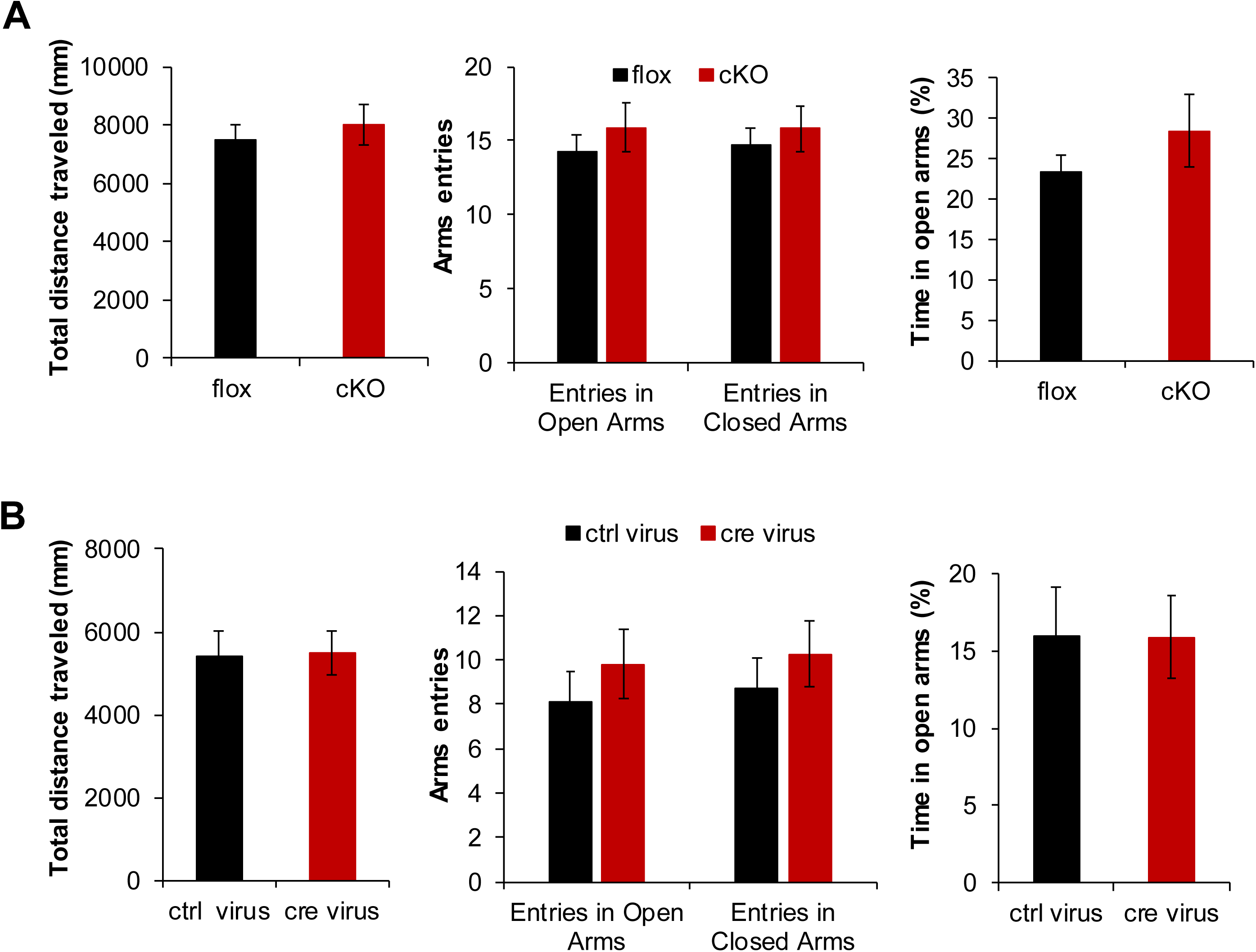
Neither Camk2α-cre^+/-^::Rapgef2^cko/cko^ (cKO) mice nor Rapgef2^cko/cko^ (flox) mice with AAV9-hSyn-cre-eGFP (cre virus) injection in the NAc showed changes in anxiety and locomotor activity, compared to their corresponding controls, in the zero maze test. All animals were housed in divided cages before behavioral experiments.

**Extended Data Figure 6-2.**
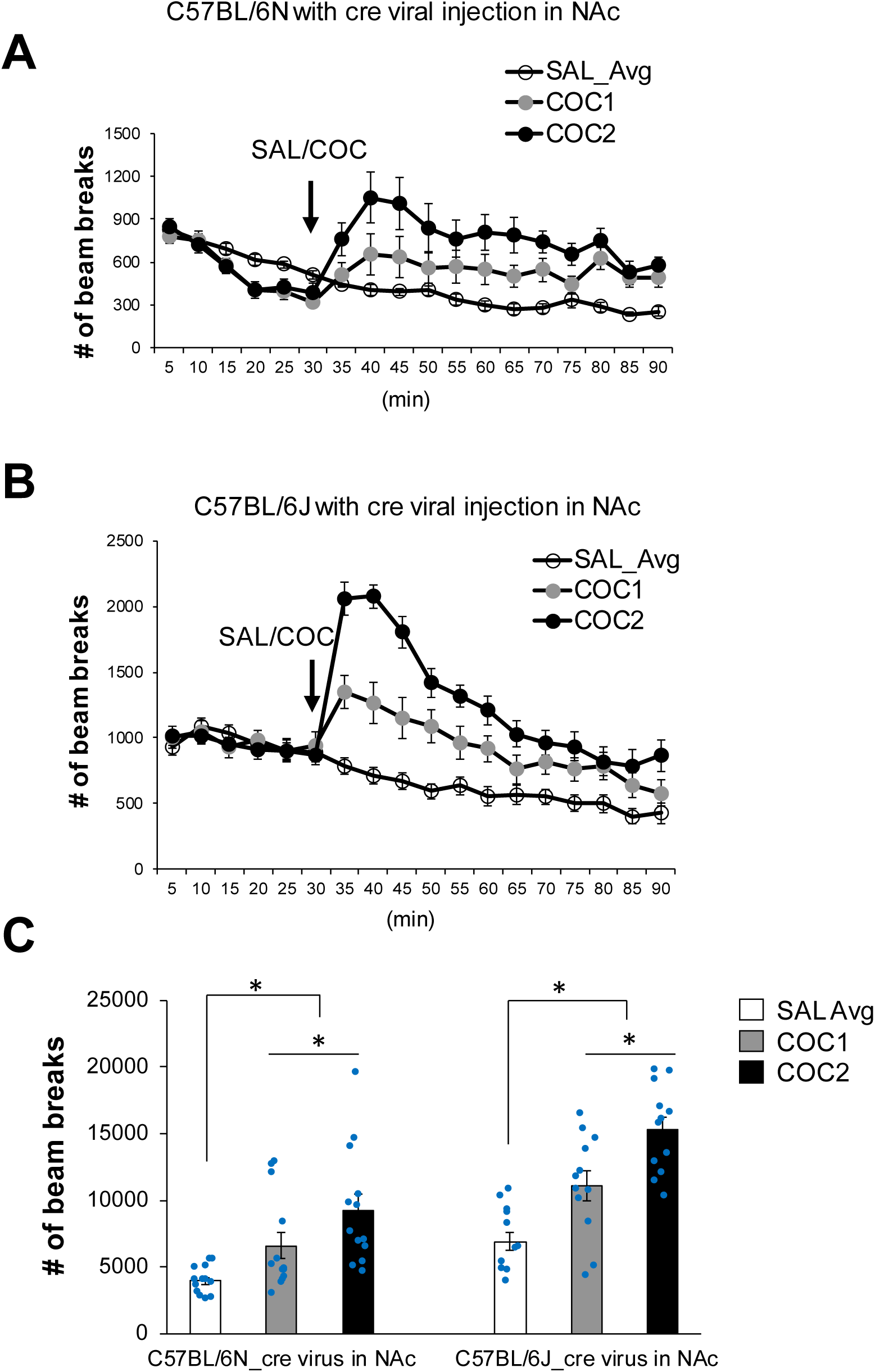
Locomotor sensitization can be induced by cocaine in C57BL/6 wild-type mice with cre viral infusion in the NAc bilaterally. Same two-injection protocol as figure 6 was used. C57BL/6N (n=13) or C57BL/6J (n=12) were injected with AAV9-hSyn-cre-eGFP bilaterally (0.3 μl of virus/each side, ∼0.5×10^9^ infectious particle per microliter). Four weeks later, animals were subjected to locomotor sensitization task. C57BL/6J (panel B) showed muchstronger response to cocaine than C57BL/6N (panel A). (C) Locomotor activity (total number of beam breaks) for one hour followingsaline or cocaine administration. C57BL/6J with cre virus injection in the NAc showed similar locomotor sensitization in response to cocaine (15 mg/kg) as Rapgef2^cko/cko^ mice or Rapgef2^cko/cko^ injected with control virus indicated in figure 6. Two-way ANOVA followed by *post hoc* Fisher’s LSD test, *p<0.05. (C57BL/6J vs C57BL/6N: F _(1, 69)_ =34.928, p<0.001; SAL Avg *vs* COC1 *vs* COC2: F _(2, 69)_ =27.006, p<0.001; genotype X drug interaction: F _(2, 69)_ =1.337, p=0.269.).

## Notes

### Competing Interest Statement

The authors have declared no competing interest.

